# Evolved differences in *cis* and *trans* regulation between the maternal and zygotic mRNA complements in the *Drosophila* embryo

**DOI:** 10.1101/737536

**Authors:** Emily L. Cartwright, Susan E. Lott

**Author notes:** Corresponding Author Information: Emily L. Cartwright, Department of Evolution and Ecology, 2320 Storer Hall, University of California, Davis, Davis, CA 95616, Susan E. Lott, Department of Evolution and Ecology, 2320 Storer Hall, University of California, Davis Davis, CA 95616. Data availability: The raw sequencing reads and processed data are available at GEO/NCBI at accession number GSE136646.

## Abstract

How gene expression can evolve depends on the mechanisms driving gene expression. Gene expression is controlled in different ways in different developmental stages; here we ask whether different developmental stages show different patterns of regulatory evolution. To explore the mode of regulatory evolution, we used the early stages of embryonic development controlled by two different genomes, that of the mother and that of the zygote. During embryogenesis in all animals, initial developmental processes are driven entirely by maternally provided gene products deposited into the oocyte. The zygotic genome is activated later, when developmental control is handed off from maternal gene products to the zygote during the maternal-to-zygotic transition. Using hybrid crosses between sister species of *Drosophila* (*D*. simulans, *D. sechellia*, and *D. mauritiana*) and transcriptomics, we find that the regulation of maternal transcript deposition and zygotic transcription evolve through different mechanisms. We find that patterns of transcript level inheritance in hybrids, relative to parental species, differ between maternal and zygotic transcripts, and maternal transcript levels are more likely to be conserved. Changes in transcript levels occur predominantly through differences in *trans* regulation for maternal genes, while changes in zygotic transcription occur through a combination of both *cis* and *trans* regulatory changes. Differences in the underlying regulatory landscape in the mother and the zygote are likely the primary determinants for how maternal and zygotic transcripts evolve.

## INTRODUCTION

Since the proposal that regulation of gene expression may be one of the primary drivers of morphological evolution was introduced (King and Wilson 1975; Carroll 1995), much research has been directed toward elucidating mechanisms underlying the evolution of gene expression. The mechanistic basis of regulatory control determines the substrate for evolution of gene expression. At the transcriptional level, gene expression is controlled by *cis*-regulatory elements (regulatory DNA sequences, such as enhancers or promoters) and by *trans* acting factors (such as transcription factors or miRNAs that bind the regulatory DNA of many genes). Due to their modular structure (Britten and Davidson 1969; Davidson and Peter 2015), changes in *cis*-regulatory elements can lead to the evolution of altered gene expression and thus new traits (Prud’homme *et al*. 2006; Chan *et al*. 2010; Kvon *et al*. 2016) without the pleiotropic consequences for other critical traits controlled by the same gene. On the other hand, transcription factors that bind to enhancers often have higher pleiotropy and thus can affect the expression of many genes. This poses a fundamental mechanistic question as to how gene regulatory evolution occurs: whether changes are more likely to occur in *cis*-regulatory elements or *trans-*acting factors.

In order to determine the relative contributions of *cis* and *trans* elements to the evolution of gene expression genome-wide, previous studies implemented the use of genetic hybrids and methods of detecting allele-specific expression (Wittkopp *et al*. 2004; Landry *et al*. 2005; Graze *et al*. 2009; McManus *et al*. 2010; Coolon *et al*. 2014; León-Novelo *et al*. 2014). Several studies point to differences in *cis* regulation as the primary mechanism of change in transcript abundance within or between species (Graze *et al*. 2009; Mack *et al*. 2016), while other studies indicate that *trans* changes are more widespread (McManus *et al*. 2010; Glaser-Schmitt *et al*. 2018). Previous research has proposed that the difference in the abundance of *cis* versus *trans* changes affecting gene expression can be explained by the divergence times of the strains or species being compared (Coolon *et al*. 2014), or the particular tissue type examined (Buchberger *et al*. 2019). Given this body of previous work, it is surprising that, to our knowledge, no previous study has compared *cis* and *trans* contributions to gene expression evolution across developmental stages in a model organism.

In this study, we ask whether contributions of *cis* and *trans* changes to gene regulatory evolution differ across developmental stages. We chose stages during embryogenesis that are close together in developmental time, yet are likely to broadly differ in the mechanistic basis of regulation. The general regulatory architecture of a particular stage likely affects how regulation at this stage can evolve. The early stages of development utilized here are under regulatory control of entirely different genomes: that of the mother, and that of the zygote. The earliest developmental processes in embryogenesis are regulated by maternally provided RNA and protein, which lay the foundation for the rest of development (Tadros and Lipshitz 2009; Vastenhouw *et al*. 2019). These maternally derived gene products carry out all initial developmental events because at the time of fertilization, the zygotic genome is transcriptionally silent. Because the zygotic genome is not yet transcriptionally active, post-transcriptional mechanisms also play an important role in regulating the amount of maternal gene products present (Tadros *et al*. 2007; Rouget *et al*. 2010; Barckmann and Simonelig 2013). As the zygotic genome is activated, control of developmental processes is handed off from the maternally deposited factors to those derived from the zygotic genome in a process known as the maternal-to-zygotic transition (MZT). The MZT is a highly conserved and regulated process during early development that occurs in all animals and in some species of flowering plants (Baroux *et al*. 2008; Tadros and Lipshitz 2009; Vastenhouw *et al*. 2019).

Regulation of early zygotic gene expression has been extensively studied over many years (Mannervik 2014; Schulz and Harrison 2019), while regulatory control of maternal genes is not well understood. Studies of maternal transcripts have largely focused on their transport into the oocyte (Mische *et al*. 2007; Kugler and Lasko 2009), their localization and movement within the oocyte and embryo (Theurkauf and Hazelrigg 1998; Kugler and Lasko 2009), their post-transcriptional regulation (Salles *et al*. 1994), and their degradation (Tadros *et al*. 2007; Bushati *et al*. 2008; Laver *et al*. 2015). Zygotic gene expression is carefully controlled, with classic examples such as the *even-skipped* gene having multiple enhancer elements along with multiple transcription factors responsible for producing complex expression patterns in developmental time and embryonic space (Small *et al*. 1992; Perry *et al*. 2011; Mannervik 2014). In contrast, maternal transcripts are produced by support cells called nurse cells during oogenesis, which are polyploid and highly transcriptionally active (Kugler and Lasko 2009; Lasko 2012), rapidly producing large amounts of transcripts. Roughly 50-75% of the genome is maternally deposited (Tadros *et al*. 2007; De Renzis *et al*. 2007; Thomsen *et al*. 2010; Lott *et al*. 2011; Vastenhouw *et al*. 2019) and there is considerable post-transcriptional regulation of maternal factors (Tadros *et al*. 2007; Bushati *et al*. 2008; Laver *et al*. 2015). For these reasons, regulation at the transcript level may not need to be as precise. While the mechanisms behind maternal transcription are not well understood, the regulatory environments driving the production of the maternal and zygotic transcriptomes are quite different.

To study the regulatory basis of differences in transcript levels at the maternal and zygotic stages of early development between species, we focused on three closely related species of *Drosophila* (*D. simulans, D. sechellia*, and *D. mauritiana*). Despite having a relatively close divergence time of 250,000 years (McDermott and Kliman 2008), these sister species have differences in the pools of transcripts present in the developing embryo both at a stage where only maternal transcripts are present, and at a stage after zygotic genome activation (ZGA; Atallah and Lott 2018). By comparing hybrids and parental lines of the species *D. simulans, D. sechellia*, and *D. mauritiana*, we asked whether, at each of these two developmental stages, changes in gene expression between species occurred due to changes in *cis*, in *trans*, or in a combination of the two.

We found that patterns of gene regulatory changes between species are distinct across developmental stages for maternally deposited transcripts and for genes with primarily zygotic expression (see Methods) in early embryogenesis. Differences in maternal transcripts occur much more frequently due to *trans* as opposed to *cis* regulatory changes, while differences in zygotic gene transcription occur through a mix of *cis, trans*, and the combined action of *cis* and *trans* regulatory changes. The complex pattern of changes found in our study at the zygotic stage speaks to what has been known about regulation at this stage for some time, that both *cis* and *trans* elements are necessary for the intricate control of gene expression in time and space at this stage of embryogenesis. The large proportion of *trans* regulatory signal found at the maternal stage may reveal fundamental properties of the regulatory architecture during oogenesis. *Trans* regulators can affect a large number of genes at the same time, as might be necessary to maximize mRNA production to load sufficient numbers of transcripts into the egg. We also identified motifs associated with *trans* regulation at the level of chromatin at the maternal stage, which lends evidence to an emerging picture of how gene expression might be regulated during oogenesis. Overall, we find distinct patterns of gene regulatory changes at the two embryonic timepoints, before and after ZGA, indicating evolved changes in gene regulation differ based on the developmental context.

## MATERIALS AND METHODS

### Crossing scheme and sample acquisition

Three *Drosophila* species were used for this study: *D. sechellia* (Dsec/wild-type;14021-0248.25) and *D. simulans* (Dsim/w[501]; 14021-0251.011) from the 12 Genomes study (Clark *et al*. 2007) and *D. mauritiana* (Dmau/[w1];14021-0241.60). For interspecific crosses, each vial was set up using 7-12 virgin females from one species and 7-10 males from another. We did not cross *D. sechellia* females and *D. simulans* males, as this combination is known to be incompatible (Lachaise *et al*. 1986). Two types of hybrid crosses were established from which embryos were collected: 1) to determine the regulatory basis of changes in zygotic gene expression, hybrid F1 embryos were collected; and 2) to determine the regulatory basis of changes in maternal gene expression, embryos produced by hybrid F1 mothers were collected. To investigate the regulatory basis of changes in zygotic gene expression, we collected hybrid F1 embryos at the very end of blastoderm stage, stage 5 (Bownes’ stages; Bownes 1975; Campos-Ortega and Hartenstein 2013), a timepoint after the zygotic genome is activated. We define late stage 5 by morphology; it is the point when cellularization is complete, but gastrulation has not yet begun. In order to determine the regulatory basis of changes in maternal gene expression, similar crosses were established with hybrid females from the F1 generations of the initial crosses and males that were the same species as the maternal species in the parental cross. We set up crosses in this manner in order to establish consistency amongst crosses, although the male genotype is unlikely to affect our data. The contribution of sperm mRNA to the zygote is debated but known contributions are small (Fischer *et al*. 2012) and likely not detectable via RNA-sequencing (Ali-Murthy *et al*. 2013). This second set of crosses was used to collect stage 2 embryos (Bownes’ stages; Bownes 1975; Campos-Ortega and Hartenstein 2013), during which time only maternal gene products are present. At this point in development, the cytoplasm has retracted from the vitelline membrane at the anterior and posterior poles of the embryo but pole cells have not yet migrated to the posterior (Ashburner 1989). As is conventional in *Drosophila* genetics, we denote our crosses by listing the female genotype first and the male genotype second. For example, in a cross between *D. mauritiana* and *D. simulans*, we write the genotype of the resulting stage 5 hybrid embryo as *mau* x *sim*. We describe the hybrid genotype of stage 2 embryos from backcrosses of hybrid F1 females and males of the maternal species in the initial cross in a similar way, e.g. (*mau* x *sim*) x *mau* (also see Figure 1 for cross diagram).

**Figure 1:**
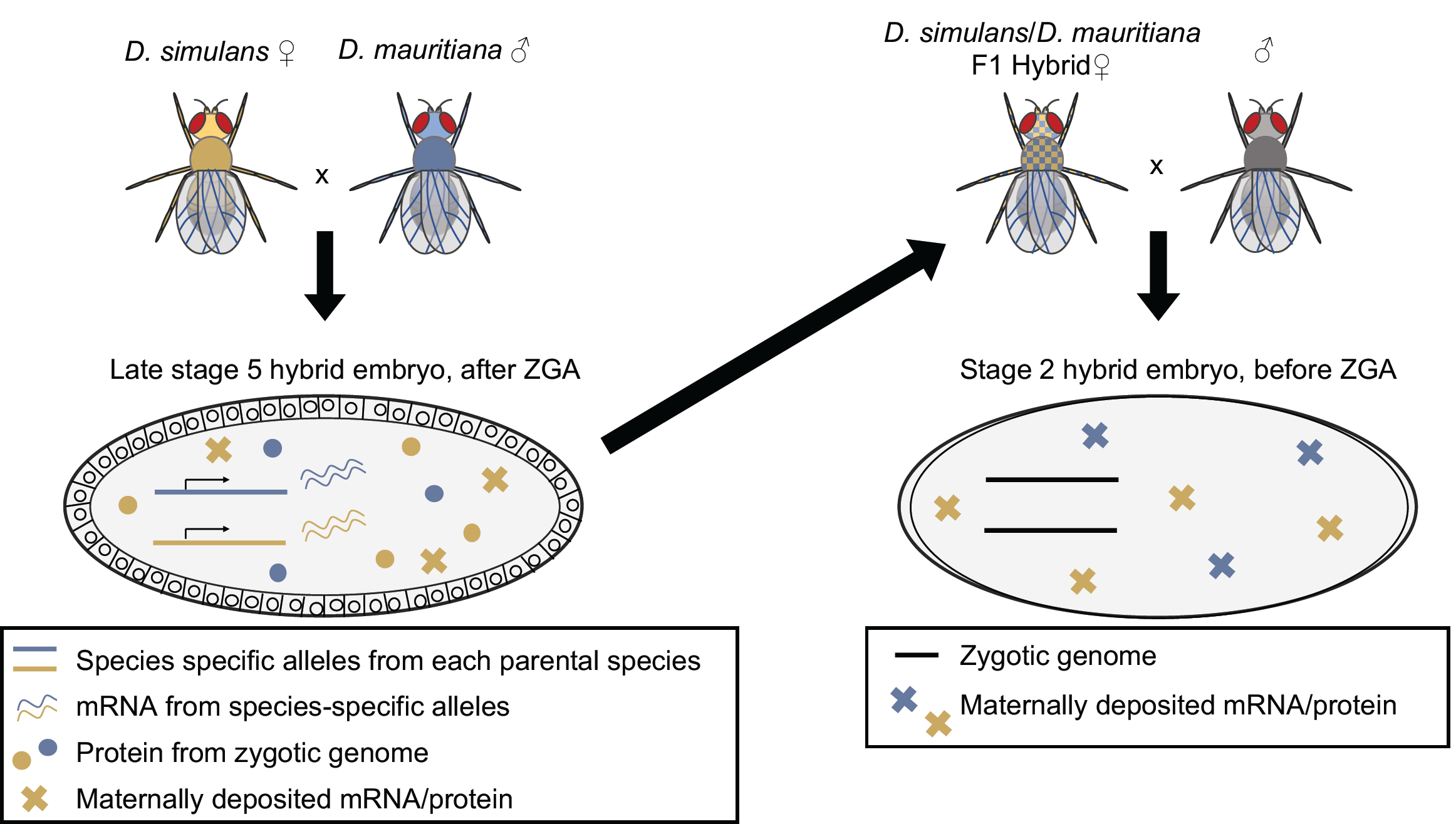
Crosses to produce hybrid embryos for the zygotic and maternal stages. To look at changes in regulation for zygotic genes, hybrid stage 5 embryos (left) were produced by crossing two parental species and collecting their eggs at the appropriate stage (late stage 5). To look at regulatory changes in maternal transcript deposition, F1 hybrid mothers were were mated to males and stage 2 embryos were collected (right). In both cases, transcription is coming from a F1 hybrid genome, either that of the zygote (left) which is measured after zygotic genome activation (late stage 5) or the mother (right) which is measured when all the transcripts in the embryo are maternally deposited (stage 2).

All flies were raised in vials on standard cornmeal media at 25°C. Flies were allowed to lay eggs for ∼2 hours (for collecting stage 2 embryos) and ∼3 hours (for collecting stage 5 embryos) before they were transferred to a new vial so that the eggs could be harvested. Eggs were collected from 4-14 day old females, dechorionated using 50% bleach and moved into halocarbon oil on a microscope slide for staging. Embryos were staged at the appropriate developmental time point under a microscope (Zeiss AxioImager M2), imaged, and promptly collected at stage 2 or at the end of stage 5 (Bownes’ stages; Bownes 1975; Campos-Ortega and Hartenstein 2013), of embryonic development.

To collect the samples after staging, the embryos were quickly transferred with a paintbrush to Parafilm (Bemis) and rolled (to remove excess halocarbon oil) into a drop of TRIzol (Ambion). The embryos were ruptured with a needle so that the contents dissolved in the TRIzol and were transferred to a tube to be frozen at -80°C until extraction. RNA was extracted using glycogen as a carrier (as per manufacturer instructions) in a total volume of 1mL TRIzol. Approximately 80-120ng total RNA was extracted from individual embryos, measured using a Qubit 2.0 fluorimeter (Invitrogen). The quality of the RNA was validated on RNA Pico Chips using an Agilent Bioanalyzer.

Genotyping was performed to determine embryo sex for stage 5 samples, as dosage compensation is not complete and transcript levels for genes on the X chromosome may differ for males and females at this time in development (Lott *et al*. 2014). DNA was extracted from each sample along with the RNA as per manufacturer instructions and amplified using a whole genome amplification kit (illustra GenomePhi v2, GE Healthcare). Sex-specific primers (Table S1) designed for use with all three species, two sets for the Y chromosome (to the *ORY* and *kl2* genes) and one control set (to *ftz*), were used to genotype the single embryos after genome amplification. For the stage 5 samples, a total of three male and three female embryos from each cross were used for sequencing. One noted exception is that in the *sim* x *mau* cross, a total of four female and two male embryos were collected, as determined from the transcriptomic data. A total of three stage 2 embryos were collected. We did not perform genotyping for embryo sex on the stage 2 embryos because the zygotic genome is not yet active at this stage in development.

### Library preparation and sequencing

The RNA from single embryos was treated with DNase (TurboDNA-free, Life AM1907) using manufacturer instructions and RNA sequencing libraries were constructed with Illumina TruSeq v2 kits following the manufacturer’s low sample protocol. The Illumina protocol uses oligo (dT) beads to enrich for polyadenylated transcripts. Because poly(A) tail length is important in determining many post-transcriptional processes during early development, including translational efficiency, it is important to ensure that the method used for mRNA selection does not produce a biased set of poly(A) tail lengths. Previous datasets report poly(A) length distributions for transcripts during oogenesis and early development (Lim *et al*. 2016; Eichhorn *et al*. 2016). We could not directly compare our data to previous reports, as these studies were done using *D. melanogaster*, which may have a different poly(A) tail length distribution than the species used in our analysis. However, previous studies comparing distributions of poly(A) tail lengths of all genes to poly(A) tail lengths of transcripts recovered through poly(A) selection in *D. melanogaster* have demonstrated that poly(A) selection with commonly used methods does not bias which transcripts are recovered from the total pool of transcripts present (Eichhorn *et al*. 2016). This includes studies that used the same single embryo approaches utilized here (Crofton *et al*. 2018; Atallah and Lott 2018). Therefore, it seems unlikely that poly(A) selection heavily biases the extracted RNA relative to the RNAs present at these developmental stages. cDNA libraries were quantified using a Qubit 2.0 fluorimeter (dsDNA BR Assay Kits) and the quality of the libraries were assessed on High Sensitivity DNA chips on an Agilent Bioanalyzer. The libraries were pooled (11-12 samples per lane) and sequenced (100bp, paired-end) in four lanes on an Illumina HiSeq2500 sequencer. Sequencing was done at the Vincent J. Coates Genomics Sequencing Laboratory at UC Berkeley.

### Data Processing

Raw reads were processed to remove adapter sequences and gently trimmed (PHRED Q<5; Macmanes 2014) using Cutadapt (version 1.7.1; Martin 2011). TopHat (version 2.0.13; Trapnell *et al*. 2012) was used to align reads to the *D. simulans* (version r2.02) and *D. sechellia* (version r1.3) genome assemblies (from the twelve species project, downloaded from Flybase) and to the *D. mauritiana* MS17 assembly (Nolte *et al*. 2013). Because the *D. mauritiana* line used for sequencing and the line used to construct the genome assembly differed, variant sites from the lab line, called using Genome Analysis Toolkit’s (GATK) Haplotypecaller, were incorporated into the MS17 assembly using Pseudoref (http://yangjl.com/pseudoRef/; Xu *et al*. 2020). Additionally, an updated annotation file for the MS17 assembly (Torres-Oliva *et al*. 2016) was used during alignment and in subsequent processing steps. Annotation files for *D. simulans* and *D. sechellia* were obtained from the same versions of the genome release of each species. Read alignment, mismatches, edit distance, and gap length were all set to three when using TopHat (version 2.0.13; Trapnell *et al*. 2012) to allow for a higher rate of read alignment.

In order to differentiate reads derived from each parental species, variant sites were called between the genomes of the species used in this analysis. RNA-seq reads from parental species samples (from previous data from Atallah and Lott, 2018, GEO accession GSE112858) were aligned to every other parental genome in each pairwise comparison using TopHat (version 2.0.13; Trapnell *et al*. 2012). The BAM files from the TopHat output were sorted and indexed using Samtools (version 1.2; Li *et al*. 2009). Picard tools (version 2.7.1) and GATK tools (Van der Auwera *et al*. 2013) were then used to identify variant sites by using the following programs: AddorReplaceReadGroups, MarkDuplicates, ReorderSam, SplitNCigarReads, and HaplotypeCaller. Additionally, indels were excluded and sites with single variants selected using the SelectVariants tool. The variants were ordered using a Pysam script (Python version 2.7.10) and read assignments to parental genomes were subsequently organized with custom R scripts using the variant sites that exist between the parental genomes (R version 3.4.1; R Core Team 2017) (Files S1, S2, S3 and S4). This pipeline was also used to update the *D. mauritiana* MS17 assembly (Nolte *et al*. 2013) with variants present in the line we used in the lab (Dmau/[w1];14021-0241.60). Normalization to the upper quantile was performed across all samples in each set of pairwise species comparisons. This was used to account for differences in the number of reads for each sample as a result of sequencing.

Read counts represent the number of reads mapping to variant sites within a gene. A cutoff of 5 or more reads mapped to any given gene was set to determine if a gene was expressed. Genes with read counts <5 in both species in any pairwise comparison were not considered to be expressed in either species so were removed from the analysis. This cutoff was tested empirically and was set to exclude genes with low count numbers that had a higher frequency of mapping in a biased manner to both parental genomes. Genes analyzed in this analysis were also limited to those with annotated orthologs in both species in any pairwise comparison. An orthologs table from Atallah and Lott, 2018, was updated using the annotations available on FlyBase (v2017) and an updated set of annotations from Torres-Oliva, *et al*. 2016. This revised orthologs table (Table S2) was used to compare genes between each species and in each direction of mapping.

Mapping bias due to differing genome quality may occur when using two different reference genomes. In our study, mapping bias can result when a higher proportion of reads from one allele map to the genome of a species used in the cross than reads from the other allele. In order to alleviate mapping bias that may occur when mapping the parental and hybrid samples to each parental reference genome, Poisson Gamma (PG) models (León-Novelo *et al*. 2014) were employed to calculate mapping bias for every set of mappings, in each pairwise comparison of species. We compared the 95% confidence intervals (CIs) from PG models (with fixed bias parameter, *q* = 0.5) in each direction of mapping. We set a slightly conservative standard for classifying allelic imbalance where genes with CIs below 0.49 or above 0.51 were called differentially expressed. Genes with CIs close to 0.50 did not appear differentially expressed when looking at the count data, so we used a more conservative cutoff. Genes that appeared differentially expressed in one direction of mapping (CIs fell outside of the range of 0.49 - 0.51 when comparing the expression levels of parental alleles in each replicate) but not in the other direction of mapping were removed from the analysis, as this was determined to be a result of mapping bias between the two genomes. We also removed genes that had disparate confidence intervals in the two mapping directions (i.e. one mapping direction yielded a CI that fell above the 0.49 - 0.51 range and the other direction of mapping yielded a CI that fell below the 0.49 - 0.51 range).

We found that between 9.6% and 10.9% of genes expressed at stage 2 (with a count >5) mapped in a biased way to parental genomes when compared to the total number of orthologous genes between any pair of species. Each between-species comparison has a different number of orthologous genes so the proportion of biased genes varies based on the pair of species compared in a cross. In contrast to the maternal stage, we found that between 5.0% and 6.2% of genes expressed at stage 5 mapped in a biased way to parental genomes when compared to the total number of orthologous genes between a pair of species. Overall, when looking at the total proportion of biased genes, not just those that were called “expressed” in our analysis, we found that between 24.8% and 28.3% of genes at stage 2 and between 20.0% and 21.2% of genes at stage 5 mapped in a biased manner when compared to the total number of orthologous genes in any comparison between species in a cross. All the genes that mapped in a biased manner were removed from our analysis. Genes that were not biased in their mapping and had a read count of >5 reads were retained for analysis.

### Genes used for stage 5 analysis

To focus on the gene regulation from the zygotic genome after ZGA, we removed genes with high levels of maternal transcript deposition from our analysis. We limited the pool of genes analyzed to those that are mostly zygotic because roughly half of maternal transcripts are not entirely degraded by stage 5 (although studies are somewhat variable in the percent reported; Tadros *et al*. 2007; De Renzis *et al*. 2007; Thomsen *et al*. 2010; Lott *et al*. 2014) and we wanted to examine only those genes that have a larger contribution to expression from the zygotic genome. For this analysis, we included genes with “zygotic-only” expression (those that are not maternally deposited) and genes that are “mostly zygotic” (those with 8-fold higher expression at stage 5 relative to stage 2, a log_2_ difference greater than three). We added a count of 1 to the transcript count for genes at the maternal and zygotic stages when calculating the log_2_ difference to avoid errors associated with taking the log_2_ of zero. We tested several cutoffs but chose the 8-fold threshold because at this conservative cutoff, most genes with high maternal transcript deposition are removed from the analysis. Additionally, for this analysis we used confidence intervals and averages generated from only female samples for genes on the X chromosome because dosage compensation is not complete at stage 5 (Lott *et al*. 2011).

### Correlation analysis and PCA

We performed correlation analysis (Figure 2, Table S3) between single embryos across replicates, stages, and genotypes in R (R Core Team 2017) using the Spearman option within the *corr* function. Principal component analysis (PCA) was also performed in R using the *prcomp* function (Figure S1).

**Figure 2:**
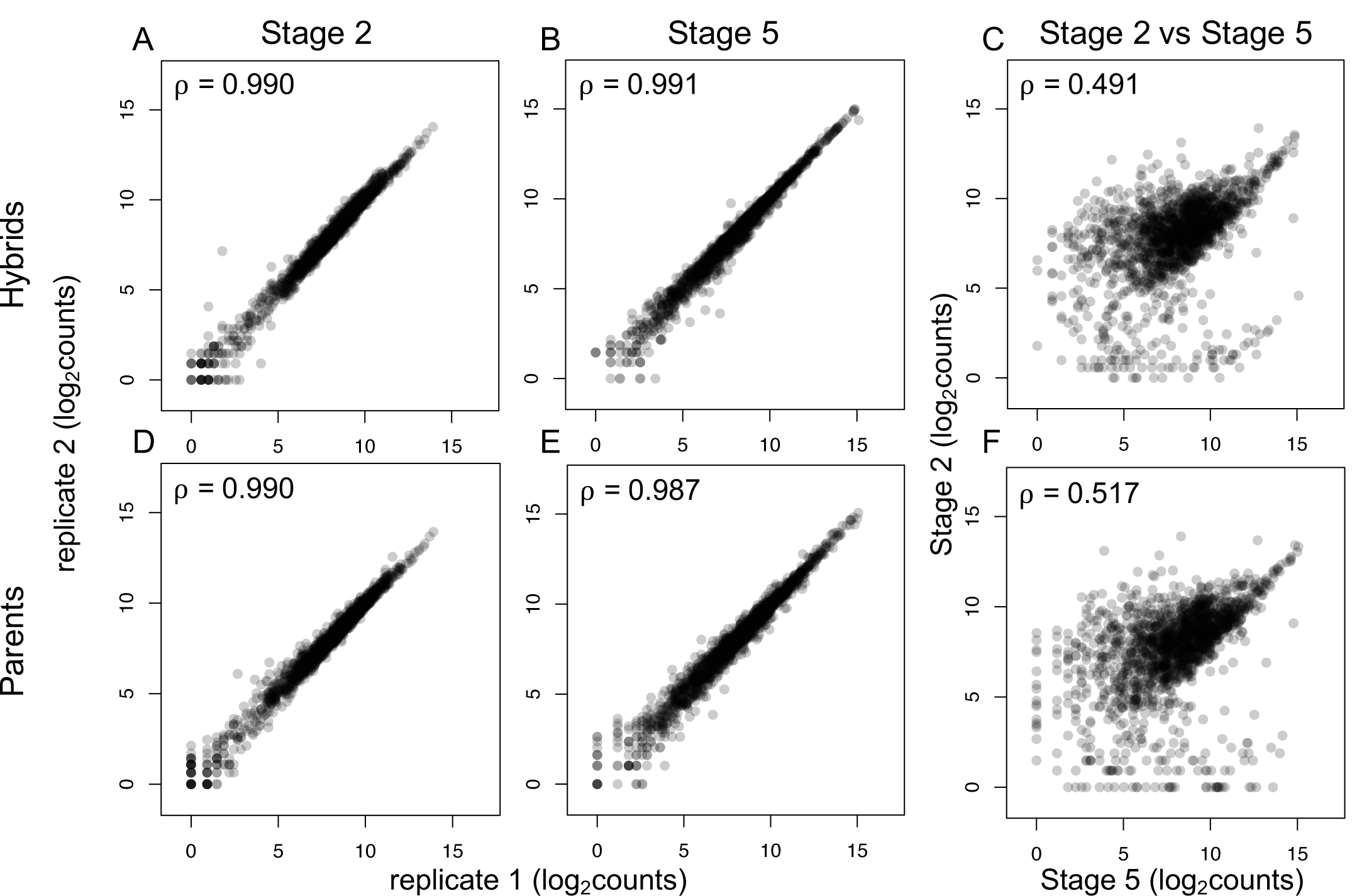
Hybrid and parental species single embryo transcript levels are highly reproducible. (A,B,D,E) Spearman’s rank correlation coefficients are high when counts from replicate transcriptomes of the same stage and genotype are compared. Correlation coefficients are similarly high in parental species (D,E) and when comparing replicates from hybrid crosses (A,B). (C,F) Samples from different stages and the same genotype have much lower correlations, indicating a large difference in transcriptomes between the maternal and zygotic stages.

### *Cis/Trans* analysis

To identify evolved regulatory changes between species, we first determined which genes showed differential expression between alleles using the 95% CIs from PG models (León-Novelo *et al*. 2014) that were also used to interpret mapping bias. We compared expression levels of alleles for individual genes within the hybrids and also between parental samples using this model. Genes with CIs that fell outside of the 49% - 51% range were defined as differentially expressed, either within the hybrid or between parental samples, while those falling within the 49% - 51% range were identified as having the same level of expression. Genes were then categorized using *cis, trans, cis* + *trans, cis x trans, compensatory*, and *conserved* categories as described in Landry, *et al*. 2005; McManus, *et al*. 2010; and Coolon, *et al*. 2014 (Figures 3, S2 and S3). We assigned the following categories for regulatory change based on the CIs generated from PG models for individual genes (see Figure 4 for individual examples):

**Figure 3:**
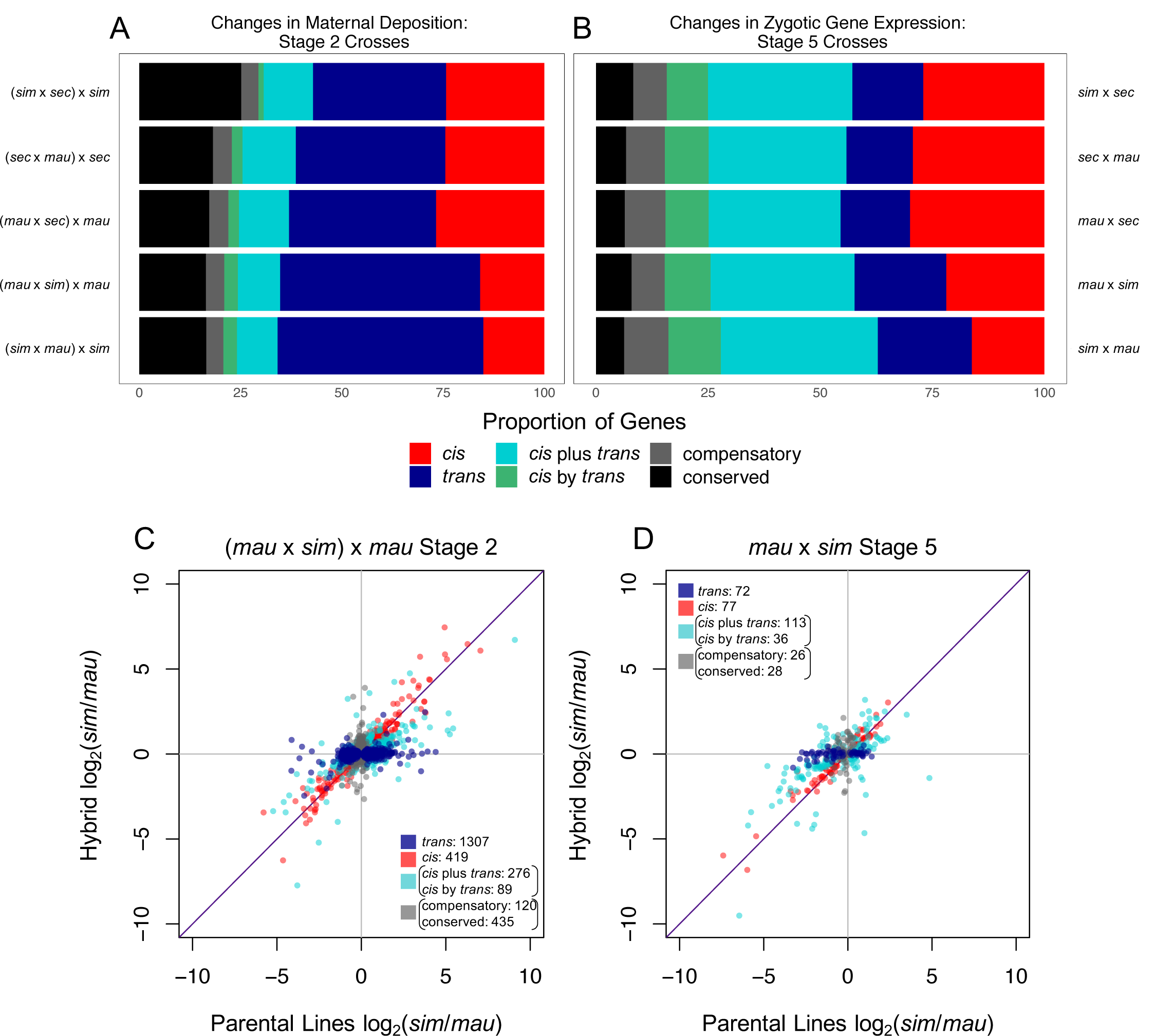
Different types of evolved regulatory changes dominate in maternal transcript deposition vs. zygotic transcription. Proportion of genes that fall into categories of regulatory change for each cross are shown for both the maternal transcript deposition (A) and zygotic gene transcription (B), for mostly-zygotic genes. Transcript level ratios between parental lines and within hybrids at stage 2 (C) and stage 5 (D) describe regulatory changes between *D. mauritiana* and *D. simulans* in one direction of crosses (for the rest of the crosses, see Figures S2 and S3).

**Figure 4:**
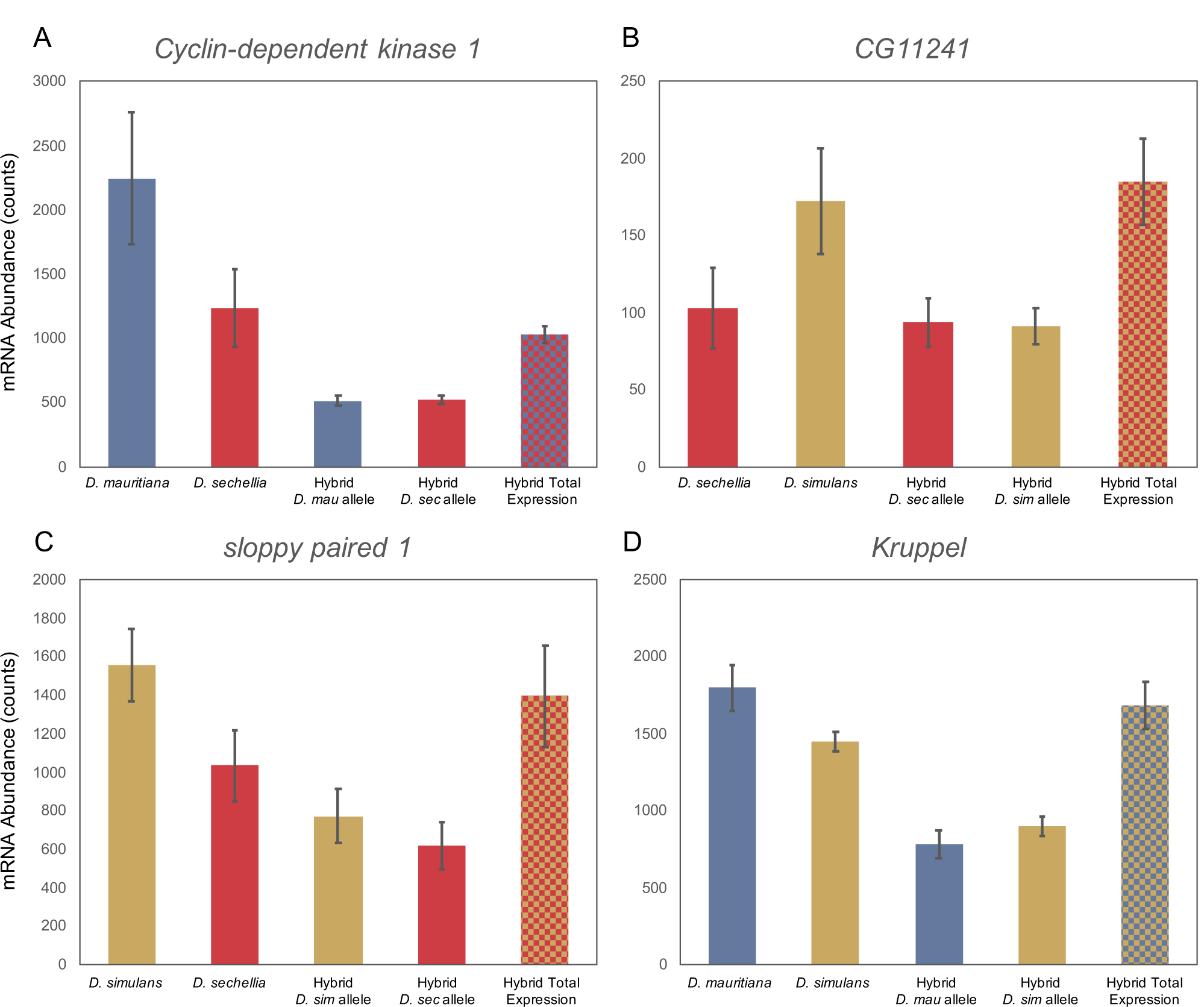
Examples of the type of regulatory changes observed, for individual genes. Transcript abundance, shown in counts, for each gene is plotted for the total mRNA abundance in both parental lines and for each parental allele within the hybrid; error bars shown represent the standard deviation. Total transcript abundance in the hybrid (the summation of levels from parental alleles in the hybrid) is shown as the last bar on the right in each graph. (A) Maternal transcript deposition of *Cyclin-dependent kinase 1* (*Cdk1*), a critical cell cycle regulator in early development, changes in *trans* regulation between *D. mauritiana* and *D. sechellia*. Hybrid mRNA abundance is from the (*mau* x *sec*) x *mau* cross. (*Cdk1* also changes in *trans* regulation in the reciprocal cross comparison, (*sec* x *mau*) x *sec*.) (B) Maternal transcript deposition of *CG11241*, a gene of currently unknown function, changes in *trans* regulation between *D. sechellia* and *D. simulans*. Hybrid mRNA abundance is from the (*sim* x *sec*) x *sim* cross. (C) At stage 5 in development, *sloppy paired 1*, a critical pair-rule segmentation gene, changes in regulation through a combination of *cis* and *trans* regulatory changes (*cis* + trans) between *D. simulans* and *D. sechellia*. Hybrid expression is shown for the *sim* x *sec* cross. *Sloppy paired 2* also changes in *cis* regulation between these two species. (D) At stage 5, *Kruppel*, a gap gene crucial to segmentation changes in regulation through a combination of *cis* and *trans* regulatory elements (*cis* x *trans*) between *D. mauritiana* and *D. simulans*. Here, expression in the hybrid is from the *sim* x *mau* cross but in the reciprocal cross (*mau* x *sim*), *Kruppel* also changes in *cis* x *trans* regulation.

#### cis

Genes categorized as having changes in *cis* are those that are differentially expressed (CIs do not overlap 49% - 51%) between the parental species and in the hybrids. (CIs for parental species and hybrids overlap each other for changes purely in *cis*. To determine this, we used the CIs generated from mapping to the *D. simulans* genome for *D. simulans/D. mauritiana* and *D. simulans/ D. sechellia* comparisons and CIs generated from mapping to the *D. sechellia* genome for *D. sechellia/D. mauritiana* comparisons.)

#### trans

Genes that are differentially expressed between the parental species (CI does not overlap 49% - 51%) but are not differentially expressed in the hybrid (CI overlaps 49% - 51%).

#### cis + trans

Genes that are differentially expressed in the hybrids and between the parental species (CI does not overlap 0.49% - 0.51%) and the CI is in the same direction for both the parents and the hybrid (i.e. both are greater than 51% but the CIs for the parents and hybrid do not overlap. For this comparison, we used the CIs generated from mapping to the *D. simulans* genome for *D. simulans/D. mauritiana* and *D. simulans/ D. sechellia* comparisons and CIs generated from mapping to the *D. sechellia* genome for *D. sechellia/D. mauritiana* comparisons.)

#### cis x trans

Genes that are differentially expressed in the hybrids and between the parental species (CI does not overlap 49% - 51%) and the CI is in opposite directions for the parents and the hybrid (i.e. one is greater than 51%, the other is less than 49%)

#### compensatory

Genes that are not differentially expressed between the parental species (CI overlaps 49% - 51%) but are differentially expressed in the hybrids (CI does not overlap 49% - 51%).

#### conserved

Genes are not differentially expressed between the parental species or within the hybrids (CIs overlap 49% - 51%).

### Inheritance Patterns

Previous studies from Gibson, *et al*. 2004 and McManus, *et al*. 2010 identified and outlined ways to classify inheritance patterns of transcript abundance in hybrids in relation to parental samples. We used these methods in our study to compare the averages of total expression levels in the hybrids relative to those of parental samples. Gene expression was considered conserved if the expression level between parental samples and the total expression in the hybrid (sum of the expression of the two species-specific alleles in the hybrid) were within 1.25-fold of one another, a log_2_-fold change of 0.32. Overdominant genes were expressed at least 1.25-fold more in the hybrid than in either parent while underdominant genes were expressed at least 1.25-fold lower in the hybrid than in either parent. Genes that were expressed at an intermediate level in the hybrid in comparison to the parental species samples involved in the cross were defined as additive. Dominance was determined when the hybrid had expression within 1.25-fold of one of the parental species such that total transcript levels in the hybrid was more similar to transcript levels in one parental species than in the other parental species.

### Candidate transcription factor identification

We took a computational approach to identify potential transcription factors that may change in *trans* regulation between the species in our analysis. We used motif enrichment programs to find potential binding sites in the upstream regions of genes changing in regulation in *D. sechellia* and *D. simulans*. We omitted *D. mauritiana* from this analysis because the *D. mauritiana* genome is not as well annotated as the genomes for *D. simulans* and *D. sechellia*. We used the Differential Enrichment mode in MEME (Bailey and Elkan 1994) as well as the findMotifs.pl script in HOMER (Heinz *et al*. 2010) to identify overrepresented motifs in the regions 500bp upstream of the annotated starting location for genes changing in regulation or with conserved regulation between species in every set of comparisons at stage 2. We utilized a 500bp region as this was empirically determined to give the highest enrichment of signal for motifs in target genes relative to background (see Omura and Lott 2020 for more information). In MEME, we used options to find motifs with any number of repetitions and a motif width of 8-12. We used default options for HOMER and supplied a background FASTA file for enrichment analysis. The background lists supplied were 500bp upstream regions from all annotated genes in the species except for those that were in the target set (either those genes with conserved or changing regulation in any set of comparisons). The 500bp regions were extracted from FASTA files (versions were the same as ones used for mapping) for each species using BEDTools (Quinlan and Hall 2010). Significantly overrepresented motifs in the target lists relative to the background supplied were then compared against databases of known transcription factor binding sites using Tomtom (MEME suite) and HOMER. All enriched motifs that appeared in both HOMER and MEME analyses are included in Table S4. All potential targets of discovered motifs with significant E-values (MEME) or high Match Rank scores in HOMER (>0.8) are also listed in Table S4 (see Figure S4 for transcript levels of differentially maternally deposited targets in embryos of parental species).

### Gene Ontology

Gene ontology (GO) analysis was done with the statistical overrepresentation test in PANTHER (Mi *et al*. 2019) using the default settings. We looked at the GO complete annotations for biological processes and molecular function but did not find any significant terms represented in the cellular component categories. For this analysis, we set a cutoff of Bonferroni adjusted p-value < 0.05. We searched for enrichment of GO categories amongst genes that change in *trans* in each cross, compared to the background of genes that are expressed (having a count >5) in each cross. We used REVIGO (Supek *et al*. 2011) to reduce the number of redundant GO categories and used the small (0.5) level of similarity as a cutoff for redundant GO terms. GO categories shared between two or more crosses at stage 5 are represented in Figure 5 and GO categories unique to a cross are shown in Figure S5. All enriched categories are listed in Table S5.

**Figure 5:**
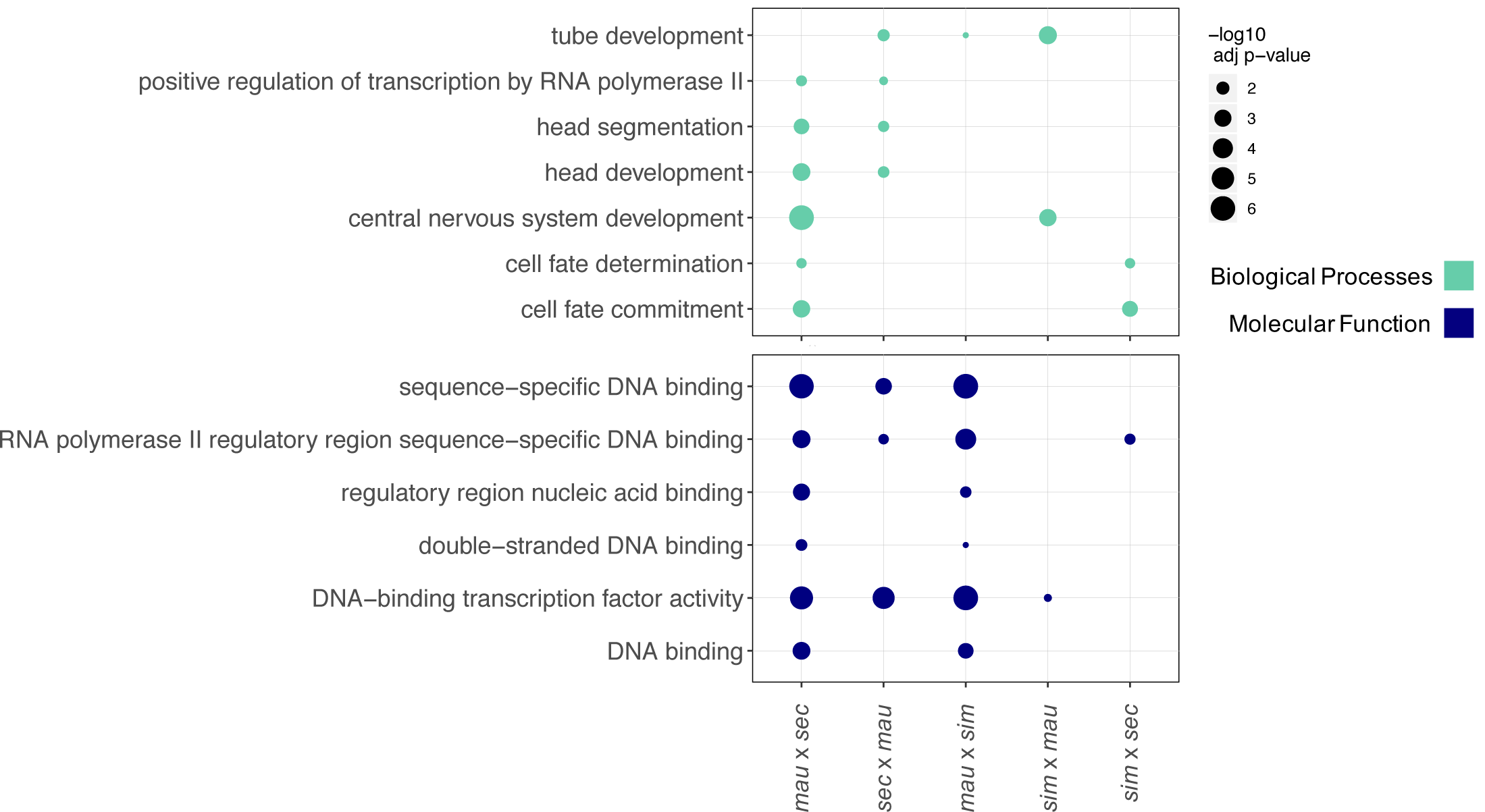
Gene ontology (GO) analysis identifies transcription factors that act in developmental processes as types of genes that change zygotically. Significantly enriched GO terms are listed for zygotically transcribed genes that change in *trans* regulation between each pair of species compared. Genes represented in this analysis are categorized as mostly zygotic (see Methods). Terms are listed for Biological Processes and Molecular Function categories and only terms that appear in more than one cross are shown in this figure. Terms unique to a specific cross are listed in Figure S5. Biological process categories identified relate to development, molecular function categories identify functions consistent with DNA binding and regulation.

### Data Availability

All sequencing data and processed data files from this study are available at NCBI/GEO at accession number: GSE136646.

## RESULTS

In order to compare the regulatory basis of evolved changes in gene expression at different stages of early embryogenesis, one stage where all the transcripts are maternally provided and the other after zygotic genome activation (ZGA), we performed a series of crosses between closely related species followed by RNA-seq on resulting embryos (Figure 1). We used the sister species *D. simulans, D. sechellia*, and *D. mauritiana*, all of which may be crossed reciprocally (with the exception of *D. sechellia* females to *D. simulans* males; Lachaise *et al*. 1986). As transcripts in the two early embryonic stages of interest are produced by different genomes, that of the mother and that of the zygote, we performed crosses to produce a hybrid genome in the appropriate generation (that of the mother or that of the zygote; Figure 1). To investigate regulatory changes in zygotic gene expression, the three species were crossed pairwise (with the noted exception), to produce F1 hybrid embryos, which were collected at a stage after zygotic genome activation (end of blastoderm stage or the end of stage 5, Bownes’ stages; Bownes 1975; Campos-Ortega and Hartenstein 2013). While the zygotic genome is fully activated at this developmental stage, maternal transcripts are not yet entirely degraded so we limited our analysis to those genes that are expressed at a much higher level after ZGA than before the zygotic genome is activated (see Methods). To discover the regulatory basis of changes in maternal transcript deposition, the F1 females were crossed to males of the same species as the maternal species in the initial cross. Resulting embryos were collected at stage 2 (Bownes’ stages; Bownes 1975; Campos-Ortega and Hartenstein 2013), when all the transcripts in the egg are maternal in origin (Figure 1). Three replicate samples were obtained for each cross at stage 2, and since stage 5 features incomplete X chromosomal dosage compensation (Lott *et al*. 2011), 6 replicates were obtained for each cross at late stage 5 (3 female and 3 male embryos with one noted exception, see Methods). mRNA-sequencing libraries were constructed from each embryo sample using poly(A) selection. Libraries were sequenced paired-end, 100bp, on an Illumina HiSeq2500.

### Reproducibility of Single Embryo RNA Sequencing Data

Previous studies have shown that single-embryo RNA-seq data is highly reproducible, despite replicate samples representing both biological and technical replicates (Lott *et al*. 2011, 2014; Paris *et al*. 2015; Atallah and Lott 2018). Our current study extends this to include replicates of F1 crosses and subsequent backcrosses between closely related species, which are as reproducible as the within-species replicates. Spearman’s rank correlation coefficients are high between replicate samples of the same species or cross at the same developmental stage (Figure 2, A,B,D,E, Table S3). For example, when comparing the RNA-seq data from stage 2 samples of the (*mau* x *sim*) x *mau* and (*sim* x *mau*) x *sim* hybrid crosses, correlation coefficients range from 0.965 to 0.995 (Table S3). Stage 5 hybrids from the *mau* x *sim* cross have equally high correlation coefficients, ranging from 0.980 to 0.996 (Table S3). Similarly, correlation coefficients for the RNA-seq data from *D. simulans* stage 5 embryos, when compared with other *D. simulans* stage 5 embryos, range from 0.985 to 0.990. The high correlation coefficients between replicates may be due, in part, to the removal of genes with differential mapping to either parental genome and those genes with very low transcript abundances (see Methods) from this analysis.

Transcript levels for embryos of the same stage but different genotypes (parental lines and hybrids) are highly similar, as indicated by their Spearman’s rank correlation coefficients (Table S3), with one notable exception. When we compare RNA-seq profiles from stage 5 hybrids to stage 5 embryos of the paternal species in the cross, we see more divergent patterns of gene expression than when we compare stage 5 hybrids to stage 5 embryos of the maternal species in the cross. This is due to the fact that many maternal transcripts are still present at the zygotic stage, and thus the hybrid zygotic embryo has many remaining maternal transcripts from the maternal species, not the paternal species, in the cross. For example, comparisons between *D. simulans* stage 5 embryos and stage 5 embryos of the *sim* x *mau* cross, where *D. simulans* is the maternal species in the cross, yield high correlation coefficients, ranging from 0.955 to 0.972. In contrast, correlation coefficients are much lower when comparing *sim* x *mau* stage 5 hybrid embryos to stage 5 embryos of the paternal species in the cross, *D. mauritiana*, ranging from 0.863 to 0.887. In this particular comparison, the lower correlation coefficients are likely due to having *D. simulans* as the maternal species in the hybrid cross for the *sim x mau* embryos. Remaining maternal transcripts are from the *D. simulans* alleles and likely explain why these hybrid embryos correlate more highly with *D. simulans* stage 5 embryos.

In contrast to highly correlated samples within a stage, comparing the transcript abundance of different stages yields strikingly lower correlation coefficients (Figure 2C and F, Table S3), emphasizing the turnover of transcripts between these stages. For example, when comparing stage 2 hybrids from crosses with *D. mauritiana* and *D. simulans* to stage 5 hybrids from the same cross, correlation coefficients range from 0.483 to 0.573 (Table S3). The correlation coefficients are lower when comparing transcript abundances of embryos of different stages than when comparing embryos within a stage, indicating that the pool of transcripts present at the maternal stage is different from that at the zygotic stage of development.

The above finding is reinforced by principal component analysis (PCA; Figure S1) for RNA expression profiles of the samples in each set of pairwise comparisons between species. We found that the first principal component corresponds to developmental stage and explains between 80.65% and 81.86% of the variance in the three sets of comparisons. The second principal component of this PCA accounts for between 6.94-8.44% of the variance in the three sets of pairwise comparisons between species and corresponds to genotype. This indicates that there is a more substantial difference between pools of transcripts at the maternal and the zygotic stage of development than there is between the pools of transcripts present in embryos of different genotypes (parental species and hybrids) at the same developmental stage.

### Regulatory changes at the maternal stage of development

Changes in gene expression can occur at many levels of regulation: transcriptional, post-transcriptional, translational or post-translational. Here, we address whether at the transcriptional level, changes in gene expression between species occurred due to changes in *cis* or in *trans* and whether the pattern of regulatory changes differs based on developmental stage. Changes in *cis* regulation can occur through changes in the DNA of regulatory regions proximal to the gene that they regulate. These types of regulatory changes have an allele-specific effect on gene expression. In contrast, changes in *trans* regulation typically occur via changes in factors that bind to the DNA, such as transcription factor proteins. Changes in *trans* regulation affect the expression of both alleles.

In order to determine regulatory changes in *cis* and in *trans* that lead to differences in maternal transcript deposition between *D. simulans, D. sechellia*, and *D. mauritiana*, we used Poisson Gamma (PG) models (León-Novelo *et al*. 2014). This allowed us to determine mapping bias as well as differential expression between alleles, between parental lines and within hybrid embryos (see Methods). We identified differential maternal transcript deposition between the parental lines as well as between species-specific alleles in the stage 2 embryos produced by hybrid mothers. We then compared the two sets of analyses to determine the proportion of *cis* and *trans* regulatory changes underlying differential maternal transcript deposition between species. We used the logic of Landry, *et al*. 2005 to classify genes as having changed in *cis* or in *trans* regulation, by comparing confidence intervals of the bias parameters generated through the PG models (see Methods).

We found that most regulatory changes underlying differentially maternally deposited transcripts occurred in *trans* between each pair of species examined (Figure 3A and C, Figure S2), where a change in a transcription factor or other *trans*-acting regulatory element affects both alleles equally (shown in Figure 4A and B). In all pairs of comparisons between species, the proportion of *trans* changes was higher than any other category of changes. Comparisons between *D. simulans* and *D. mauritiana* had the highest percentage of *trans*-only regulatory changes (between 49.4% and 50.8%) while comparisons between *D. simulans* and *D. sechellia* had a lower percentage of regulatory changes solely in *trans* (32.9%). The second highest proportion (between 15.0% and 26.7%) of regulatory changes between species at the maternal stage occurred only in *cis* regulation. Slightly fewer regulatory changes occurred due to a combination of *cis* and *trans* acting factors (between 13.4% and 15.8% in all comparisons). Most genes that change in *cis* and in *trans* regulation are assigned to the *cis* + *trans* category, which indicates that the allele with higher expression in the parental lines is also preserved as the allele with higher expression in the hybrid (the changes in *cis* and in *trans* affect gene expression in the same direction; Landry *et al*. 2005; McManus *et al*. 2010; Coolon *et al*. 2014). We found a smaller proportion of genes changed in regulation through *cis* x *trans* interactions, where changes in *cis* and in *trans* have opposing effects on gene expression and the allele with the lower level of expression in the hybrid is from the parental line with the higher level of expression (Landry *et al*. 2005; McManus *et al*. 2010; Coolon *et al*. 2014). We also found a percentage of genes with conserved levels of maternal transcript deposition between species, between 16.4% and 25.1% in all crosses. *D. simulans* and *D. sechellia* have the highest percentage of conserved genes while *D. simulans* and *D. mauritiana* have the lowest percentage of conserved genes. We also found a small proportion of genes, between 4.2% and 4.7% in all comparisons, that have evolved compensatory mechanisms of regulation, where the genes are not differentially expressed between the parental samples but are differentially expressed in hybrids. This implies that while transcript levels are the same between species, regulatory changes have occurred, which then become visible in the environment of the hybrid.

Genes with expression differences due to *trans* regulatory changes include regulators with critical functions in important processes governed by maternal gene products, such as *Cdk1*, a cell-cycle regulator necessary for the rapid cleavage cycles in early development (Farrell and O’Farrell 2014). Examples of individual genes with changes attributed to *trans* regulatory differences at the maternal stage are represented in Figure 4A and B.

### Binding sites for chromatin modifiers are enriched in the regulatory regions of maternally deposited genes

As *trans* regulatory changes can affect numerous genetic loci, we asked whether there are *trans* regulatory factors that may affect the differential deposition of a number of maternal transcripts between the species studied. For this, we identified binding sites in the predicted *cis*-regulatory regions of all differentially expressed genes and compared them to identified binding sites in the *cis*-regulatory regions of genes with conserved expression between species at the maternal stage. In the pool of genes with altered expression between species, we included not only those with differences in *trans* regulation, but also those with differences in other regulatory categories (*trans, cis, cis* + *trans, cis* x *trans*, and compensatory). This is because genes with changes in *cis* regulation may have had changes that affect the binding of the same *trans* regulators. For genes with differential expression, and separately for genes with conserved expression between species, we took a computational approach. We used both HOMER and MEME (Bailey and Elkan 1994; Heinz *et al*. 2010), to search for overrepresented motifs within 500 base pairs upstream of transcription start sites, as compared to the rest of the genome (see Methods). We used the upstream regions of genes in *D. simulans* and *D. sechellia* because the *D. mauritiana* genome is not as well annotated, as compared to the other two species in this study.

Interestingly, we found that the *cis*-regulatory regions from both genes with conserved and genes with differential transcript levels between species are enriched in motifs associated with insulator binding (Table S4). Specifically, we found that the Dref/BEAF-32 binding site (BEAF-32 and Dref bind overlapping DNA sequences; Hart *et al*. 1999) is the most significantly enriched (Table S4). These factors are annotated as insulators (Matzat and Lei 2014; Ali *et al*. 2016) and known to be associated with topologically associated domains (TADs) (Liang *et al*. 2014; Ramírez *et al*. 2018). The binding site for M1BP also appeared significantly enriched in both sets of genes that change in regulation and in ones that are conserved in regulation across species (Table S4). M1BP is involved in transcriptional regulation and RNA polymerase II pausing at the promoter of genes (Li and Gilmour 2013), which may also be associated with regulating chromatin state (Ramírez *et al*. 2018). Our findings are consistent with previous studies that identified the enrichment of binding sites for M1BP, BEAF-32 and Dref in the promoter regions of genes that are maternally deposited (Chen *et al*. 2013; Omura and Lott 2020). These binding sites have also been associated with housekeeping genes (Zabidi *et al*. 2015), and the pool of maternal transcripts is enriched with housekeeping genes (Liu *et al*. 2014). However, maternal genes that are not housekeeping genes are even more highly enriched for these binding sites (Omura and Lott 2020), thus the involvement of insulators or other chromatin regulators in maternal regulation is unlikely due only to the inclusion of housekeeping genes among maternal genes. Transcript abundance data from our study indicates that *Dref, BEAF-32*, and *M1BP* are differentially maternally deposited in several between-species comparisons (Figure S4), although in certain crosses, hybrid reads mapped in a biased way to *Dref, BEAF-32* and *M1BP*, and thus they were excluded from our regulatory analysis. As the motifs for these *trans*-acting factors are enriched in the upstream regions of all maternal genes, relative to upstream regions of all annotated genes, they are likely important regulators of transcription during oogenesis, and therefore also likely targets of regulatory evolution between species.

### Evolution of regulation for zygotically expressed genes

To determine the regulatory basis of changes in zygotic transcript abundance between species, we compared expression levels in late stage 5 parental species samples to late stage 5 hybrid samples and used PG models to identify *cis* and *trans* regulatory changes, similar to our maternal analysis (see Methods). We limited our analysis at the zygotic stage to those genes that are mostly-zygotic: zygotically expressed but not maternally deposited (zygotic-only) or expressed at the zygotic stage at an 8-fold higher level when compared to the maternal stage (we will refer to these as mostly zygotic genes, see Methods).

While we found that most gene expression changes at the maternal stage of development are due to changes in *trans* regulation between the three sister species, we see strikingly different patterns of regulatory changes after ZGA. At the zygotic stage, differences in gene expression between the three species examined occur mostly due to regulatory changes in both *cis* and *trans*, either by *cis* + *trans* or *cis* x *trans* interactions (Figure 3B and D, Figure S3). Changes in both *cis* and *trans* regulatory elements (either *cis* + *trans* or *cis* x *trans* interactions) account for expression differences in 39% to 47% of zygotic genes at stage 5 in our between-species comparisons. We also see a higher proportion of these interactions occurring in a *cis* + *trans* pattern (between 29% and 35% of all genes) as opposed to a *cis* x *trans* pattern (between 9% and 12% of all genes) of regulatory interactions. In contrast, *cis-*only and *trans-*only changes account for a smaller number of differences in gene expression levels at this stage in development. In all comparisons, we found between 15% and 21% of genes changing only in *trans* regulation. There are between 16% and 30% of genes that change only in *cis* regulation between each pair of species compared at this stage in development. Compared to the maternal stage, we found a larger proportion of genes with compensatory changes (between 7% and 10% of all genes) in gene regulation and a smaller proportion of genes that are conserved (between 6% and 8% of all genes) between each pair of species comparisons. The smaller proportion of genes with conserved transcript levels at the zygotic stage compared to the maternal stage is consistent with earlier findings showing maternal transcripts to be more highly conserved between species than zygotic transcripts (Atallah and Lott 2018). Examples of evolved changes include regulators critical to important early zygotic processes, such as gap gene *Kruppel* and pair-rule gene *sloppy paired 1*, which are required for segmentation along the anterior-posterior axis (Nüsslein-Volhard and Wieschaus 1980; Grossniklaus *et al*. 1992) (Figure 4C and D).

Transcriptional regulation at the maternal stage may be broadly determined by regulation at the level of chromatin, as evidenced in this work and by another study (Omura and Lott 2020). In contrast, regulation at the zygotic stage can be gene or pathway specific and involve transcription only in a spatially localized subset of cells (Jäckle *et al*. 1986; Johnston and Nüsslein-Volhard 1992). As such, if a *trans* regulator changed at the zygotic stage, it may affect genes involved in a specific developmental process. For these reasons, we wanted to ask if genes whose zygotic expression differed between species due to changes solely in *trans* regulation had a specific molecular function or were part of a particular biological process. We used PANTHER (Mi *et al*. 2019) to perform gene ontology (GO) analysis on genes changing only in *trans* regulation in each pairwise comparison of species at stage 5 (see Methods). Identifying GO categories over multiple crosses identifies the types of genes that evolve changes repeatedly over evolution. Shared categories were broad, and included those related to DNA binding, positive regulation of transcription by RNA polymerase II, cell fate determination, and several developmental categories (Figure 5). The range in GO categories represented, while broadly important at this developmental timepoint, demonstrate how genes changing in *trans* are distributed across developmental processes. As may be expected for zygotic genes at this stage in development, this finding suggests that changes in *trans* regulators of zygotic genes can affect a broad range of molecular and developmental processes. We also investigated biological process categories unique to each specific cross, these are represented in Figure S5 and Table S5.

### Modes of Inheritance in Hybrids

Misexpression in hybrid offspring has been used to examine regulatory incompatibilities that may contribute to speciation (Michalak 2003; Ortíz-Barrientos *et al*. 2006; Moehring *et al*. 2007; Mack *et al*. 2016). As the maternal and zygotic transcripts examined during embryogenesis showed different patterns of gene expression evolution, we also asked whether there were more hybrid incompatibilities present at one stage than the other by looking at whether these two developmental stages showed different levels of transcript misexpression. One way to identify misexpression in hybrids is to compare the inheritance of transcript levels in hybrids to transcript levels in each parental species. Here, we quantify the total transcript abundance for a gene by summing the levels of both species-specific alleles in the hybrid and comparing this level to the transcript abundance in both parental lines. We used methods developed by Gibson, *et al*., 2004 to define modes of inheritance in our hybrids; as in previous studies (Gibson *et al*. 2004; Landry *et al*. 2005; McManus *et al*. 2010; Coolon *et al*. 2014), we used a conservative fold change of 1.25 (log_2_-fold change of 0.32) to define those genes that do not change in the total transcript abundance between genotypes (conserved). Genes with transcript levels that are higher (overdominant) or lower (underdominant) in the hybrid relative to either parental species are categorized as misexpressed. The total transcript abundance in the hybrids can also be more similar to one parent versus the other. Here, we categorize the parental line with expression most similar to the hybrid as the dominant parent. Expression in the hybrid can also have a level intermediate to both parental species (additive).

Both the maternal and zygotic stages show a significant proportion of genes where one species’ allele is dominant (Figure 6, Table S6). In both stages, the species that is dominant in each set of crosses is consistent. A higher proportion of genes that are dominant have *D. simulans*-like expression (in any cross involving *D. simulans*) in comparison to the proportion that have expression more like the other parental line in the cross (see Table S6 for percentages). We found that *D. mauritiana* has the least dominance in any cross involving this species. Taken together, our findings indicate that *D. simulans* has the most dominant effect on gene expression at both developmental stages, while *D. mauritiana* has the least dominant effect, with dominance in *D. sechellia* falling between the other two species. While there has been previous work proposing relationships between the proportion of dominance and the physiology of unique species (McManus *et al*. 2010), it is difficult to determine any known factors between these three species that would predict this pattern of relative dominance (species range, effective population size, egg size/maternal investment). It is, however, interesting that while the proportion of changes in *cis* and *trans* vary considerably between stages that both stages have dominance among the largest categories of modes of inheritance, and that the relative patterns of which species are dominant is conserved.

**Figure 6:**
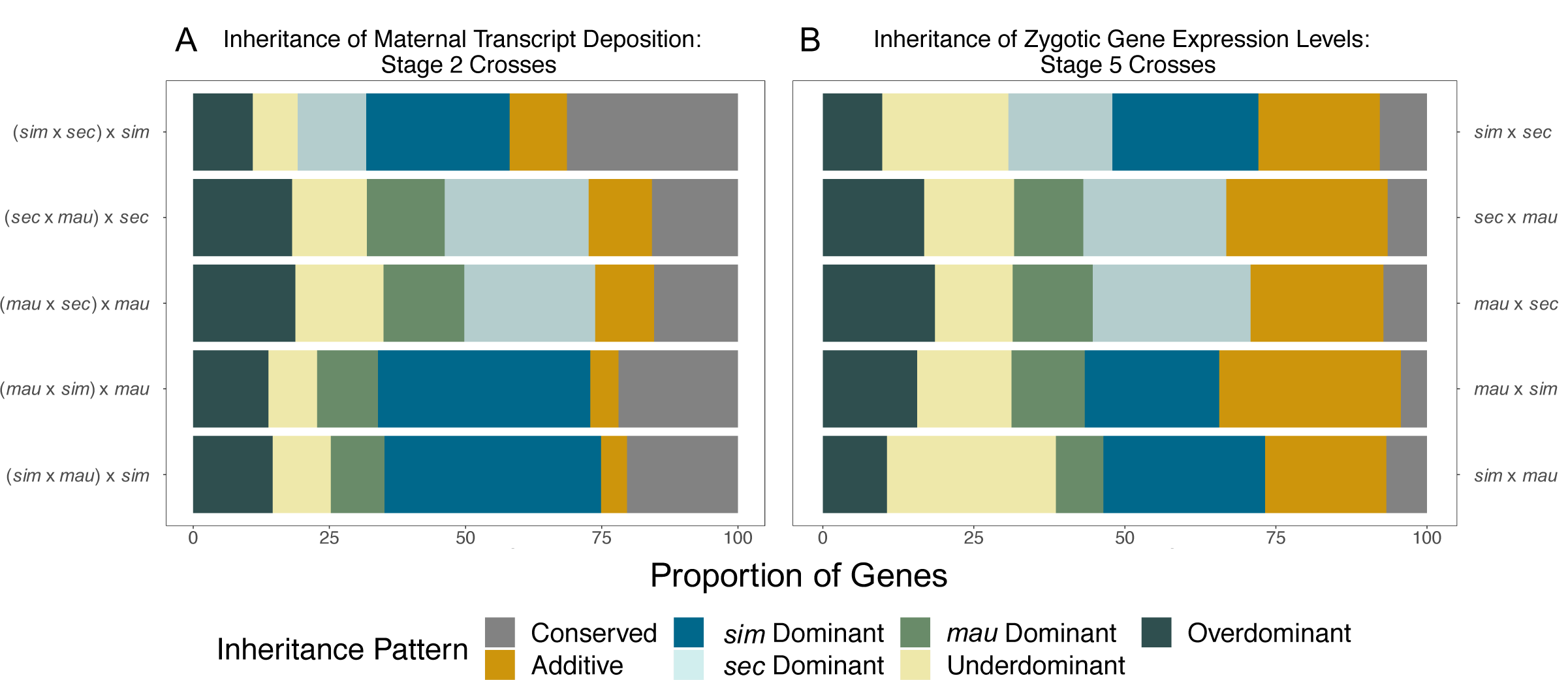
Patterns of inheritance show dominance of particular parental genomes at both stages. A) Shows patterns of inheritance for stage 2, over all genes and all crosses. B) Shows patterns of inheritance for stage 5, for mostly zygotic genes (see Methods) and all crosses. The maternal stage (A) shows a higher proportion of conserved genes than the zygotic stage (B). Both stages show a high degree of dominance for *D. simulans* for crosses involving that species, and for *D. sechellia* in crosses with *D. mauritiana*, forming the general dominance pattern of *D. simulans* > *D. sechellia* > *D. mauritiana*. There is a greater proportion of additive inheritance for the zygotic stage (B) than the maternal stage (A).

Strikingly, while many genes show conservation of expression levels between parental species and in the hybrids at both developmental stages, we found a much higher percentage of conserved transcript levels between parents and hybrids for genes that are maternally deposited (Figure 6, Table S6). We found a high proportion of genes with conserved transcript levels at stage 2 in all crosses, between 15.4% and 31.4% of all genes. In contrast, in stage 5 crosses we found conserved transcript abundance in between 4% and 8% of all genes that are either zygotic-only or are mostly zygotic (see Methods for definitions). While there is a large difference in the percentage of conserved genes between the two stages, our stage 5 analysis is limited to those genes with much higher expression at the zygotic stage in comparison to the maternal stage of development. There may be more genes that are mostly zygotic or zygotic-only that are misregulated at this stage in development relative to all of the genes that are expressed at stage 5.

In contrast to maternal expression patterns, we found more genes that have an additive mode of inheritance or that are misexpressed in the hybrids for zygotic genes (Figure 6, Table S6). Previous studies indicate that additive inheritance is associated with *cis* regulatory divergence (Lemos *et al*. 2008; McManus *et al*. 2010). This is consistent with our findings that a larger proportion of genes at the zygotic stage have expression divergence due, in part or wholly, to *cis* regulatory changes and that more zygotic genes show an additive pattern of inheritance. Higher levels of misexpression at the zygotic stage, taken together with lower conservation of transcript levels at the zygotic stage, suggests that zygotic genes may contribute more to genetic incompatibilities than maternal genes.

## DISCUSSION

In this study, we asked whether evolution of gene regulation differs at different developmental stages. We found striking differences in the proportions of *cis* and *trans* regulatory changes between the stage of embryogenesis where all transcripts are maternally derived, and a stage just a few hours later after the zygotic genome has been activated. Between the species examined, we uncovered an overwhelming number of *trans* regulatory changes resulting in differential maternal transcript levels, whereas a complex mix of *cis, trans*, and the combination of the two were responsible for changes in zygotic transcription of mostly zygotic genes (see Methods). Here, we propose that the differences in the patterns of gene regulatory evolution between the stages we examined may be due to fundamental differences in the biological context and regulatory architecture producing the transcriptomes present at these stages.

Maternal transcripts are produced by support cells called nurse cells during oogenesis and are either transported by microtubule-dependent mechanisms or dumped into the oocyte along with the cytoplasmic contents of the nurse cells upon apoptosis (Kugler and Lasko 2009). Many aspects of maternal provisioning have been well-studied in *D. melanogaster*, including transport of transcripts into the oocyte (Mische *et al*. 2007), localization of transcripts within the oocyte (Theurkauf and Hazelrigg 1998), post-transcriptional regulation of maternal gene products (Salles *et al*. 1994), and subsequent degradation of maternal transcripts (Tadros *et al*. 2007; Bushati *et al*. 2008; Laver *et al*. 2015). Surprisingly, how transcription is regulated in the nurse cells is not well understood. Nurse cells are polyploid, and are able to rapidly transcribe a large quantity of RNA that represents a large proportion of the genome (Tadros *et al*. 2007; De Renzis *et al*. 2007; Thomsen *et al*. 2010; Lott *et al*. 2011; Vastenhouw *et al*. 2019) to provide the oocyte with the large stock of transcripts needed. The oocyte itself is thought to be largely transcriptionally silent (Navarro-Costa *et al*. 2016). What we found here, and what was also found in another study investigating binding motifs in maternal factors across the *Drosophila* genus using computational methods (Omura and Lott 2020), is that maternal transcription is associated with *trans* factors annotated to be insulators and that interact with topologically associated domains (TADs). This provides evidence that maternal transcription may be controlled broadly at the level of chromatin state. In this context, we predict that changes in only a few *trans* factors can be responsible for the bulk of the between-species changes in maternal transcription. Thus, changes in the levels of *trans* regulators at this stage may easily be responsible for changes in transcription level for a number of genes.

In contrast to the large proportion of regulatory changes in *trans* at the maternal stage, differences in zygotic gene transcription for genes that are mostly zygotic (see Methods) between these species is predominantly explained by a combination of changes in *cis, trans, cis* + *trans*, and *cis* x *trans*. Zygotic gene transcription for genes without a maternal complement is fundamentally different than maternal gene transcription. Unlike the bulk transcription that takes place in the nurse cells, zygotic gene transcription is precisely regulated at the spatial and temporal level across the embryo with enhancer regions playing a large role in where, when and how much genes are expressed (Haines and Eisen 2018). Due to these fundamental differences in gene regulation, the embryo at the zygotic stage may be more sensitive to changes in gene expression than at the maternal stage. Specifically, changes in *trans* regulation, which can affect the expression of many genes, may be detrimental to the developing organism at this stage. In contrast, changes in *cis* regulation are gene-specific and may only affect gene expression in a subset of the embryo, which might be a more suitable way of fine-tuning zygotic expression. We propose that fundamental differences in the regulatory landscape and developmental purpose of the maternal versus the zygotic stage likely explain why the evolution of gene expression occurs through different mechanisms for transcripts that are maternally deposited and genes that are primarily zygotic.

While this study was directed at understanding the regulatory basis of evolution in gene expression at the maternal and zygotic stages of embryogenesis, it also provides insight into the relative conservation of gene expression, both between species and between parent and hybrid offspring. Here, in both the analysis of regulatory changes and the analysis of modes of inheritance, we found more genes with conserved transcript levels among those that are maternally deposited relative to those that are zygotically transcribed. This is in agreement with previous studies that identified high conservation of maternally deposited transcripts relative to those transcribed zygotically between species (Heyn *et al*. 2014; Atallah and Lott 2018) and indicates that the maternal stage is highly conserved. We observe lower conservation of transcript levels at the zygotic stage. A caveat our gene expression analysis at the zygotic stage is that we had to remove genes that still have a large maternal component at this stage (roughly 50% of total transcript pool at late stage 5 is maternally derived; Lott *et al*. 2014). Thus, our finding is best viewed as genes whose transcripts are primarily zygotic at stage 5, have a higher rate of evolutionary change. Additionally, the large proportion of genes with conserved transcript levels at the maternal stage may be unexpected considering that there is substantial post-transcriptional regulation of maternally deposited factors (Tadros *et al*. 2007; Rouget *et al*. 2010; Barckmann and Simonelig 2013), so it is not clear that a high degree of conservation at the transcript level should be necessary to maintaining conservation at the protein level. Alternatively, if the maternal genome is primarily regulated at the level of chromatin state, this may be a mechanistic constraint on evolution at the level of gene expression. It may be functionally difficult for a gene located in a region of open chromatin to be repressed, or for a gene in a region of heterochromatin to gain expression. Thus, it may be easier to evolve differences in expression over evolutionary time via post-transcriptional mechanisms for maternal genes. Further study is needed to disentangle conservation at the transcript and protein levels of maternal factors across species.

In addition to the differences in conservation between stages, we also found differences in the patterns of inheritance of gene expression between species at the maternal and zygotic stages of embryogenesis. The zygotic stage has a larger proportion of additive differences, which some previous theory (Gibson *et al*. 2004) and empirical studies (Lemos *et al*. 2008; McManus *et al*. 2010) have suggested may be more likely to be changes in *cis* regulation. This would be consistent both with the increased relative role of *cis* changes at the zygotic stage compared to the maternal stage found here, as well as what is known about zygotic gene regulation more generally (Mannervik 2014). In addition, a larger proportion of changes at the zygotic stage fall into the broad category of misregulation (underdominant, overdominant), which have been proposed to increase with divergence time (Coolon *et al*. 2014) and may be a potential source of hybrid incompatibility between species (Michalak 2003; Ortíz-Barrientos *et al*. 2006; Moehring *et al*. 2007; Mack *et al*. 2016).

In this study, we found that differences between species in levels of maternally deposited transcripts and zygotically transcribed genes evolve via different patterns of regulatory change. We found that maternal transcript abundance is more conserved but when changes do occur, they occur more frequently through *trans* regulation in comparison to zygotic complements. Regulatory organization, constraints, and developmental processes that are specific to each developmental stage likely play a large role in determining how gene regulation can evolve at these two embryonic timepoints. Further study is needed to characterize the molecular basis of evolved changes in transcript level on a single gene level, and more generally to determine what is controlling the regulatory landscape at each stage in development.

## ACKNOWLEDGMENTS

We would like to thank members of the Lott lab for comments on the manuscript, as well as members of the UC Davis fly community for their contributions. We would also like to thank Amanda Crofton and Anthony Le for their assistance with genotyping and RNA extraction and Graham Coop for helpful discussions. This work was supported by the National Institute of General Medical Sciences of the National Institutes of Health grant R01GM111362 and the Floyd and Mary Schwall Fellowship.

---

**Figure S1:**
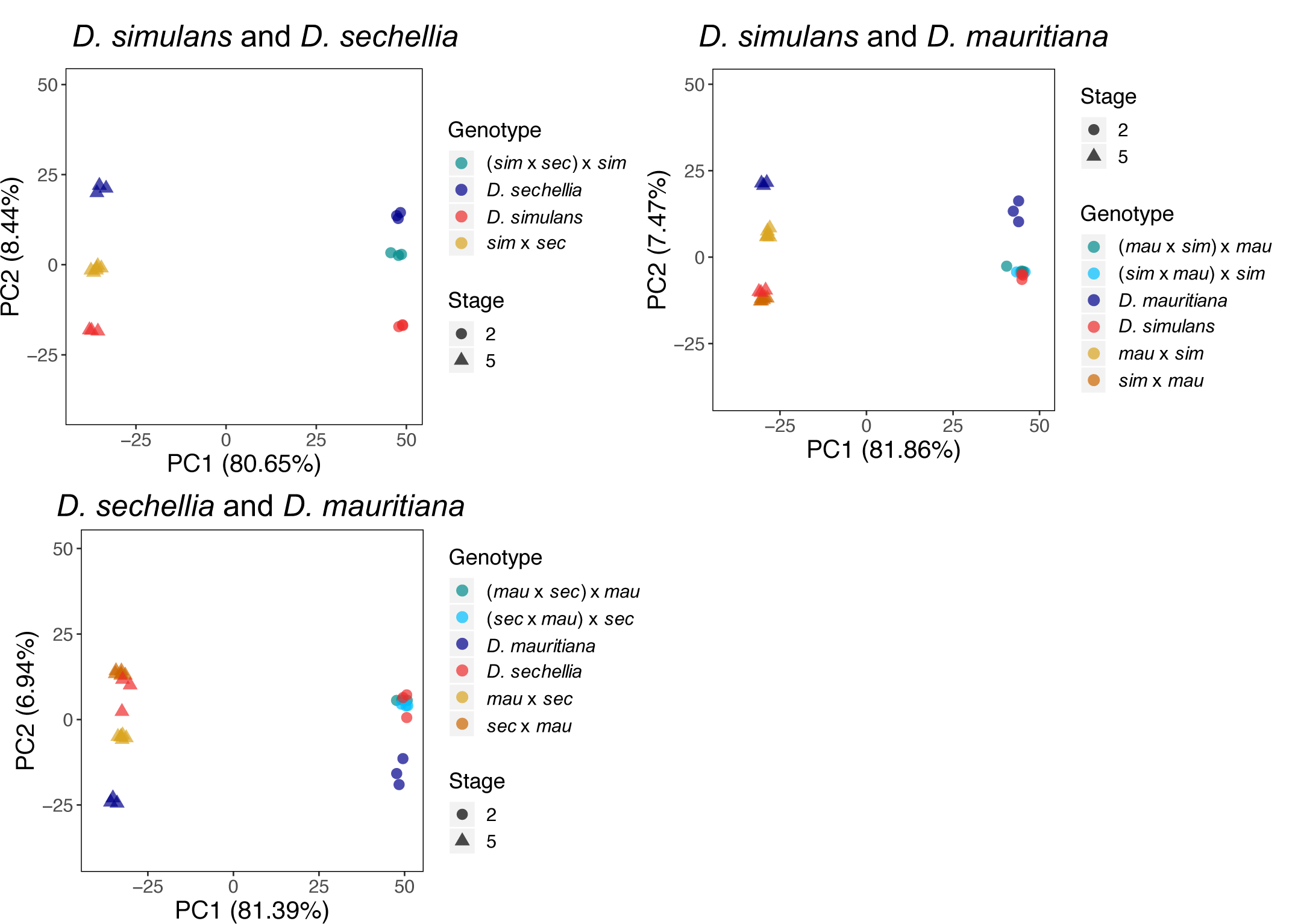
PCA plots for transcript abundance in all crosses. Samples of each stage, 2 or 5, cluster together. Samples of each genotype also cluster together, parental samples and hybrids. Proportion of variance explained by each principal component is listed on each axis.

**Figure S2:**
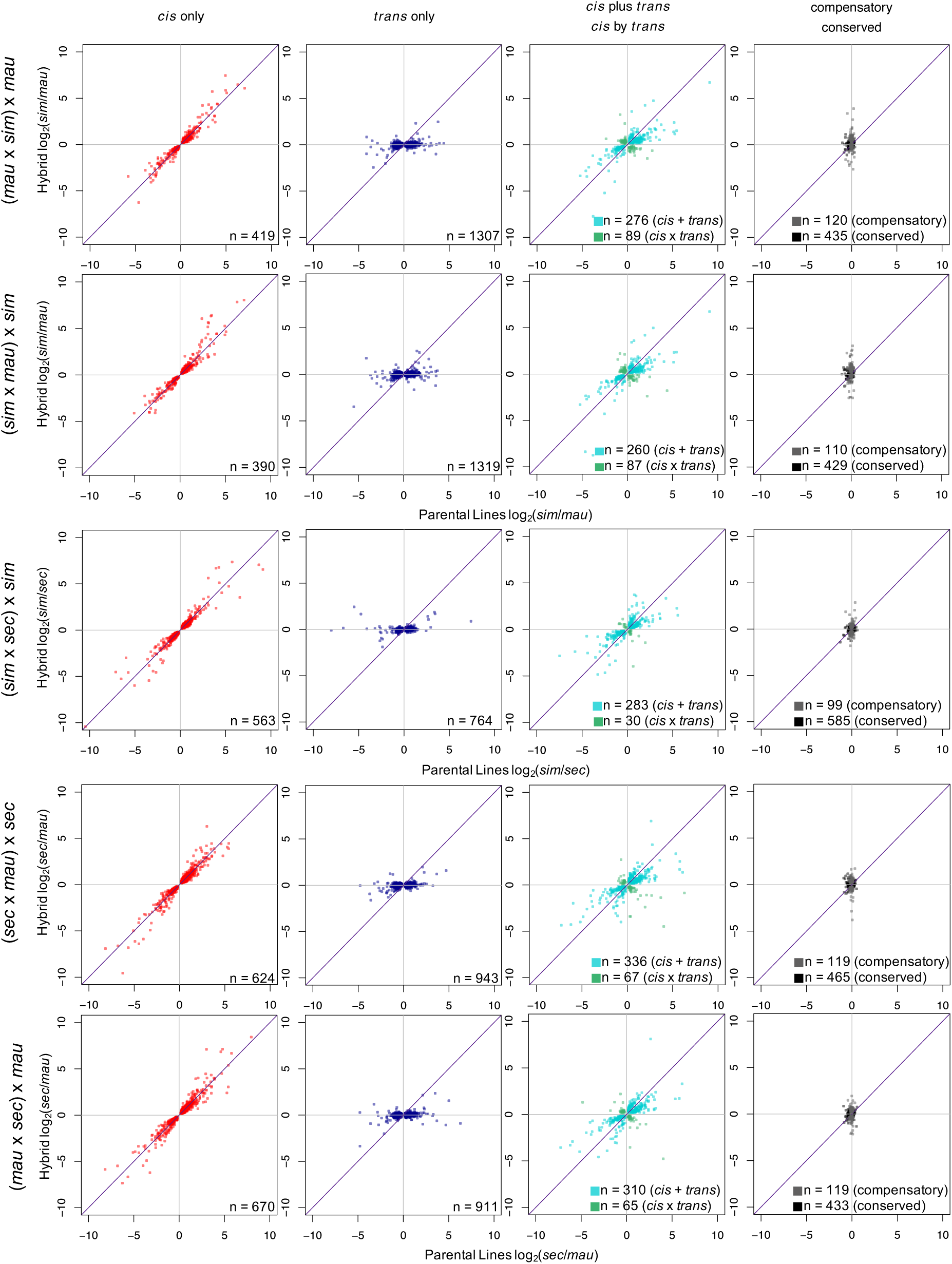
Regulatory changes in all pairwise comparisons for maternally deposited transcripts. Transcript level ratios between parental lines and within hybrids at stage 2 describe regulatory changes between species in each set of crosses. The number of genes in each category of regulatory change (n=) is listed in each plot. For definitions of categories of changes and criteria, see Methods.

**Figure S3:**
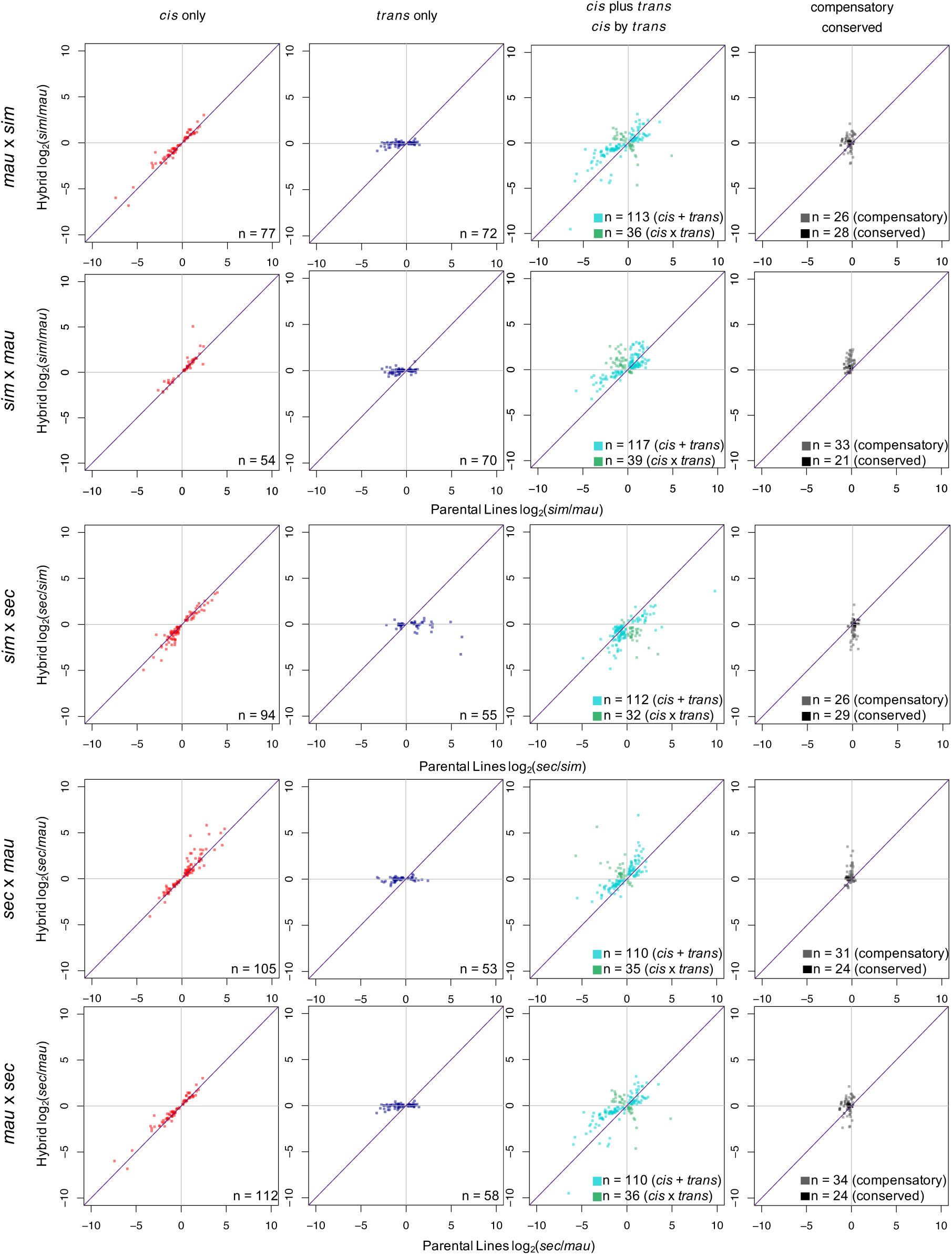
Regulatory changes in all pairwise comparisons for mostly zygotic genes. Transcript level ratios between parental lines and within hybrids at for mostly zygotic genes (see Methods) at stage 5 describe regulatory changes between species in each set of crosses. The number of genes in each category of regulatory change (n=) is listed in each plot. For definitions of categories of changes and criteria, see Methods.

**Figure S4:**
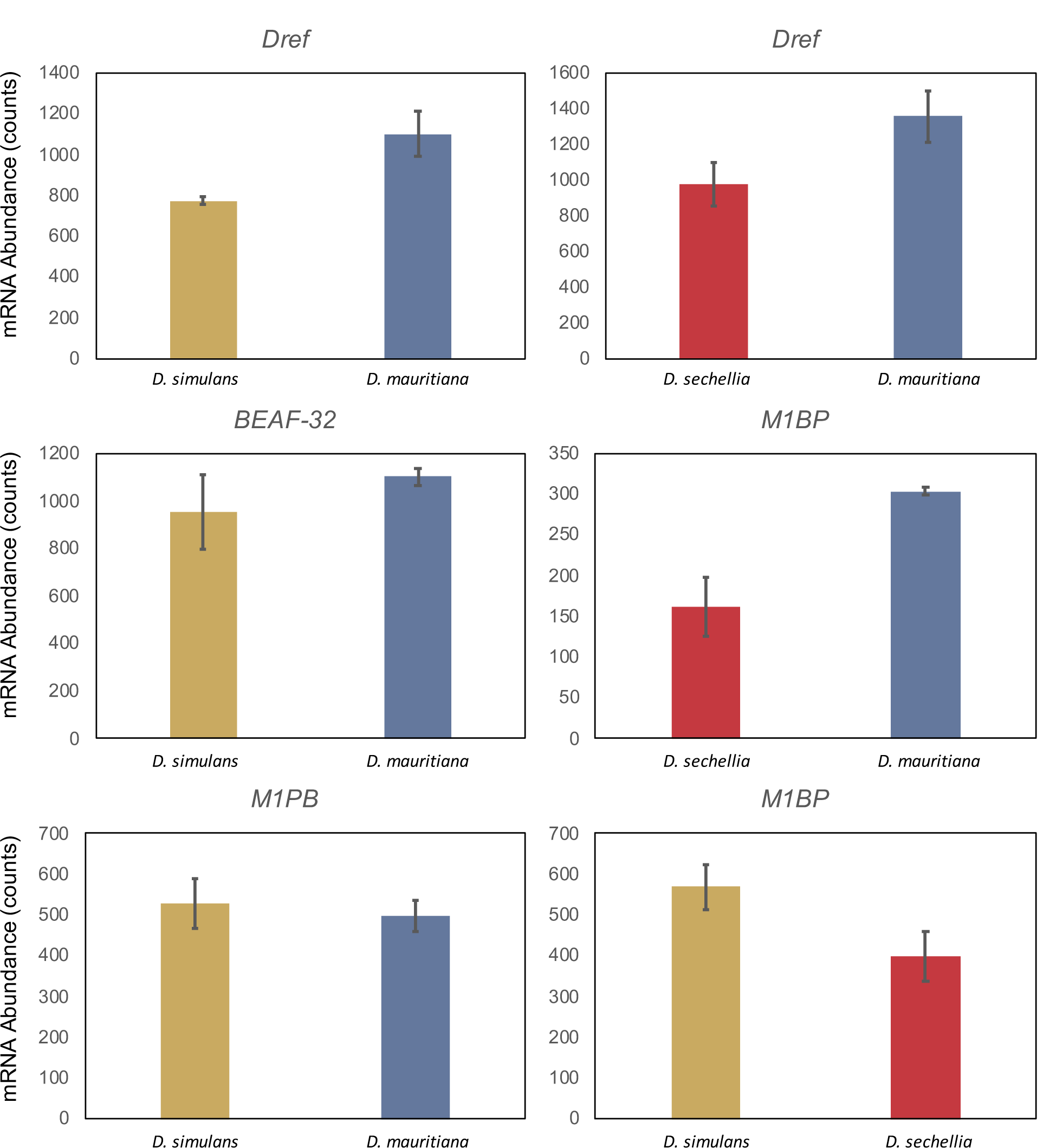
Transcript abundance from parental lines at stage 2 demonstrates differential maternal deposition of *M1BP, Dref* and *BEAF-32*. Counts for *D. simulans*/*D. mauritiana* and *D. simulans*/*D. sechellia* comparisons are averages across replicates from alignment to the *D. simulans genome*. Counts for *D. sechellia*/*D. mauritiana* comparison are averages across replicates from alignments to the *D. sechellia* genome. Error bars represent standard deviations. Count data for the same species and gene may differ across comparisons due to the genome used for alignment in each comparison and normalization of counts within a comparison.

**Figure S5:**
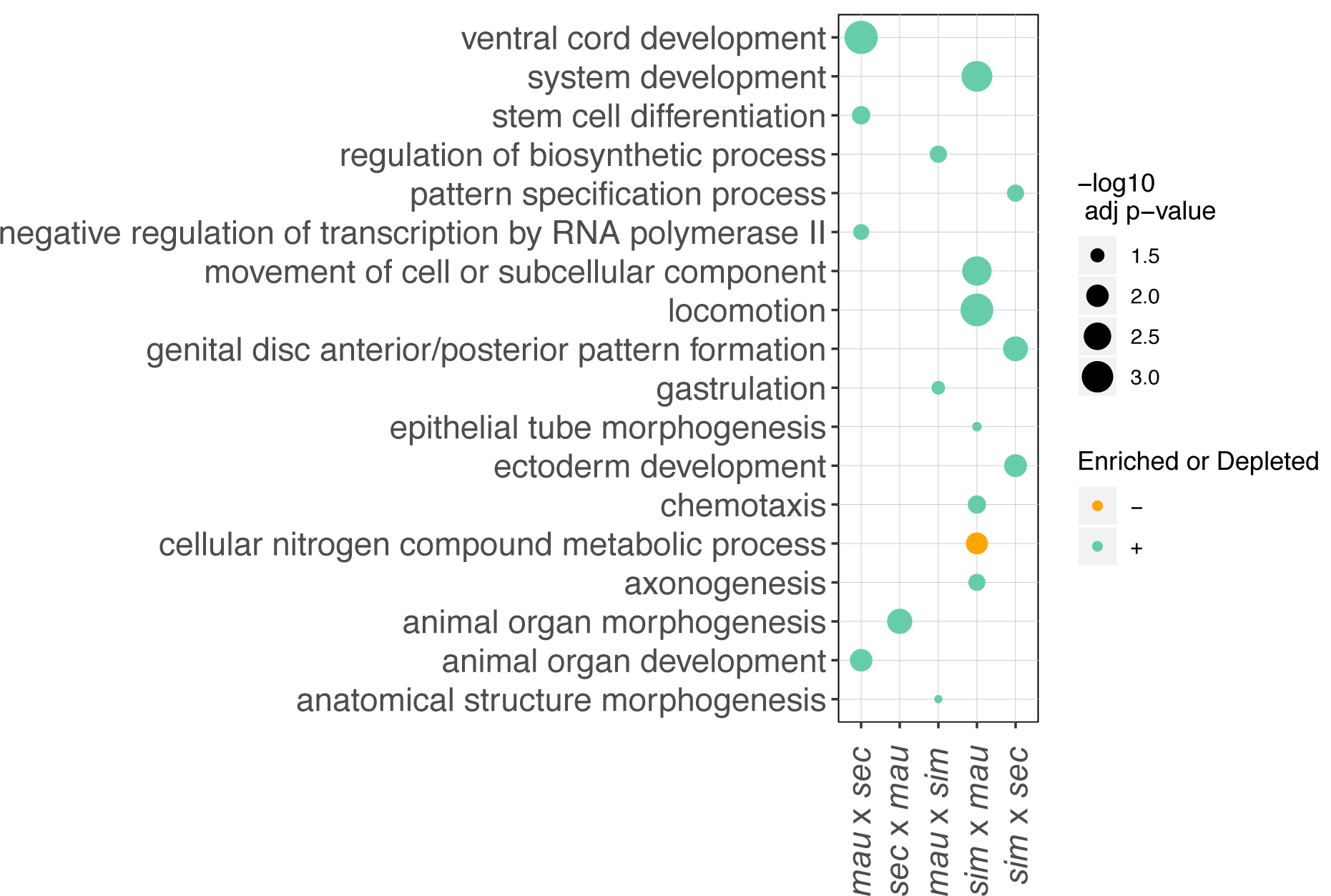
Gene ontology (GO) analysis for categories unique to a specific cross show enrichment for specific developmental processes. Significantly enriched GO terms are listed for zygotically transcribed genes that change in *trans* regulation between each pair of species compared. Again, zygotically transcribed genes are limited to those that are mostly zygotic (see Methods), in comparison to the maternal stage of development. Terms are listed for the biological processes category. Gene categories identified uniquely in a single cross primarily represent specific types of developmental processes, and may indicate evolved differences in parental genomes in these processes.

**Table S1:**
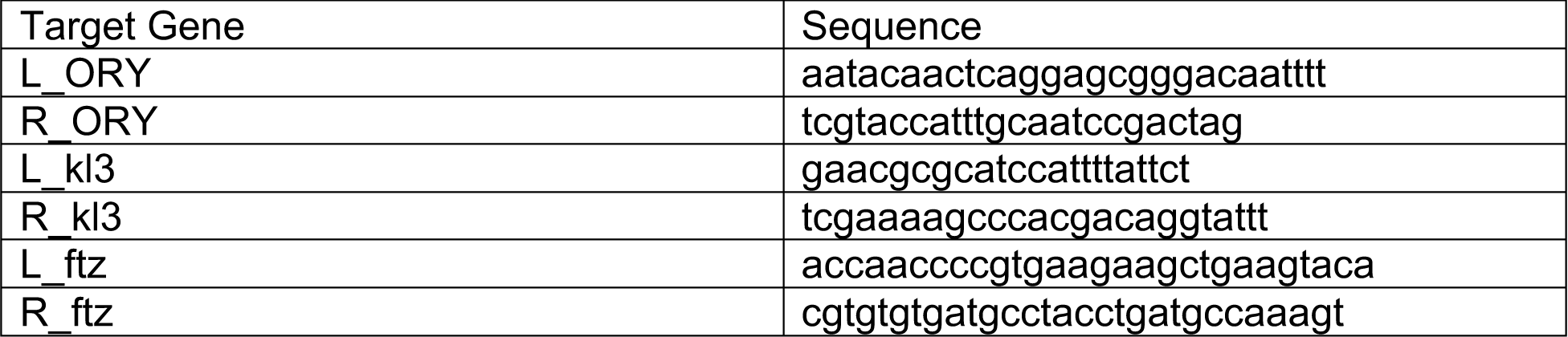
Primers for genes *ORY, kl3* (both on the Y chromosome) and *ftz* (control locus, on 3R) that were used for genotyping stage 5 embryos as male or female.

**Table S4:**
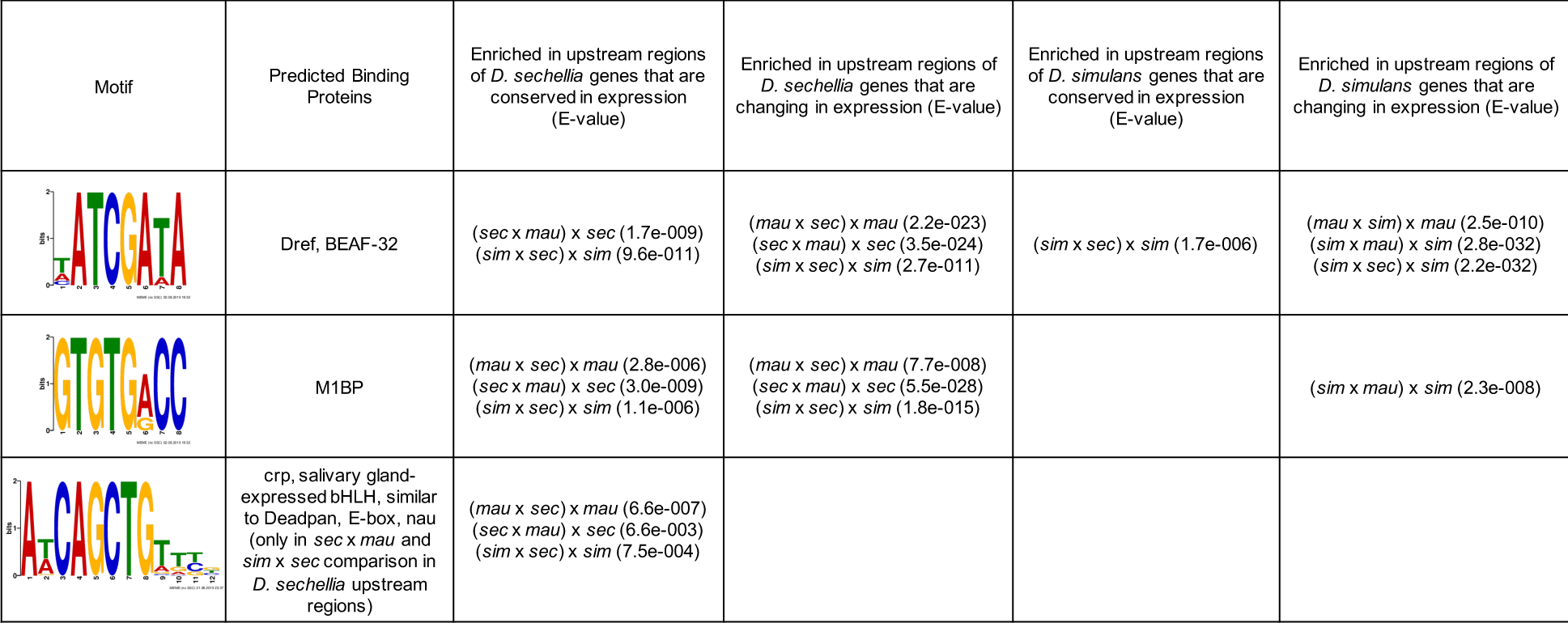
Enriched motifs found upstream of maternally deposited genes. Sequences 500bp upstream were extracted for genes in *D. simulans* and *D. sechellia* that change in regulation or that are conserved in each pairwise comparison. Motifs that were significantly enriched in analysis using MEME and HOMER are listed in the table and predicted binding proteins discovered using Tomtom and Homer are also described. E-values generated by MEME indicating the enrichment of each motif compared to background in each cross are also listed. The position weight matrix represented is a representative example of the discovered motifs.

| Motif | Predicted Binding Proteins | Enriched in upstream regions of *D. sechellia* genes that are conserved in expression (E-value) | Enriched in upstream regions of *D. sechellia* genes that are changing in expression (E-value) | Enriched in upstream regions of *D. simulans* genes that are conserved in expression (E-value) | Enriched in upstream regions of *D. simulans* genes that are changing in expression (E-value) |
| --- | --- | --- | --- | --- | --- |
|  | Dref, BEAF-32 | (*sec* x *mau*) x *sec* (1.7e-009)<br>(*sim* x *sec*) x *sim* (9.6e-011) | (*mau* x *sec*) x *mau* (2.2e-023)<br>(*sec* x *mau*) x *sec* (3.5e-024)<br>(*sim* x *sec*) x *sim* (2.7e-011) | (*sim* x *sec*) x *sim* (1.7e-006) | (*mau* x *sim*) x *mau* (2.5e-010)<br>(*sim* x *mau*) x *sim* (2.8e-032)<br>(*sim* x *sec*) x *sim* (2.2e-032) |
|  | M1BP | (*mau* x *sec*) x *mau* (2.8e-006)<br>(*sec* x *mau*) x *sec* (3.0e-009)<br>(*sim* x *sec*) x *sim* (1.1e-006) | (*mau* x *sec*) x *mau* (7.7e-008)<br>(*sec* x *mau*) x *sec* (5.5e-028)<br>(*sim* x *sec*) x *sim* (1.8e-015) |  | (*sim* x *mau*) x *sim* (2.3e-008) |
|  | crp, salivary gland-expressed bHLH, similar to Deadpan, E-box, nau (only in *sec* x *mau* and *sim* x *sec* comparison in *D. sechellia* upstream regions) | (*mau* x *sec*) x *mau* (6.6e-007)<br>(*sec* x *mau*) x *sec* (6.6e-003)<br>(*sim* x *sec*) x *sim* (7.5e-004) |  |  |  |

**Table S6:** Number of genes within each category of inheritance pattern in the hybrids. Stage 2 hybrids are shown in the first five rows and stage 5 hybrids are shown in the last five rows of the table.

| Table: Number of genes with each pattern of inheritance in hybrids |  |  |  |  |  |  |  |
| --- | --- | --- | --- | --- | --- | --- | --- |
|  | Additive | *sim*<br>dominant | *sec*<br>dominant | *mau*<br>dominant | Underdominant | Overdominant | Conserved |
| *(sim x mau) x sim* | 88 | 735 |  | 182 | 197 | 270 | 376 |
| *(mau x sim) x mau* | 97 | 733 |  | 210 | 168 | 260 | 412 |
| *(sec x mau) x sec* | 237 |  | 540 | 292 | 280 | 371 | 322 |
| *(mau x sec) x mau* | 220 |  | 489 | 302 | 329 | 382 | 313 |
| *(sim x sec) x sim* | 180 | 450 | 215 |  | 141 | 187 | 536 |
| *sim x mau* | 51 | 68 |  | 20 | 71 | 27 | 17 |
| *mau x sim* | 77 | 57 |  | 31 | 40 | 40 | 11 |
| *sec x mau* | 70 |  | 62 | 30 | 39 | 44 | 17 |
| *mau x sec* | 58 |  | 69 | 35 | 34 | 49 | 19 |
| *sim x sec* | 49 | 59 | 42 |  | 51 | 24 | 19 |

## Notes

### Competing Interest Statement

The authors have declared no competing interest.

